# Evolvability of cancer-associated genes under APOBEC3A/B selection

**DOI:** 10.1101/2023.08.27.554991

**Authors:** Joon-Hyun Song, Liliana M. Dávalos, Thomas MacCarthy, Mehdi Damaghi

## Abstract

Evolvability is an emergent hallmark of cancer that depends on intra-tumor heterogeneity and, ultimately, genetic variation. Mutations generated by APOBEC3 cytidine deaminases can contribute to genetic variation and the consequences of APOBEC activation differ depending on the stage of cancer, with the most significant impact observed during the early stages. However, how APOBEC activity shapes evolutionary patterns of genes in the host genome and differential impacts on cancer-associated and non-cancer genes remain unclear. Analyzing over 40,000 human protein-coding transcripts, we identified distinct distribution patterns of APOBEC3A/B TC motifs between cancer-related genes and controls, suggesting unique associations with cancer. Studying a bat species with many more APOBEC3 genes, we found diverse motif patterns in orthologs of cancer genes compared to controls, similar to humans and suggesting APOBEC evolution to reduce impacts on the genome rather than the converse. Simulations confirmed that APOBEC-induced heterogeneity enhances cancer evolution, shaping clonal dynamics through bimodal introduction of mutations in certain classes of genes. Our results suggest that a major consequence of the bimodal distribution of APOBEC affects greater cancer heterogeneity.

**Highlights:**

- Using a measure of the extent which genes are affected by APOBEC activity, we found that many genes are *maximally robust* in the human genome. Interestingly, the distribution in the genome of a bat that has many APOBEC genes was similar.
- In contrast, when analyzing the subset of cancer-associated genes the distribution is bimodal with many genes appearing to susceptible to APOBEC activity.
- Analysis of orthologs of human genes and other species showed a wide range in the extent to which APOBEC affects genes having the same function.
- The bimodal distribution of susceptibility to APOBEC drives greater heterogeneity in simulated clonal evolution of cancer.

## Introduction

Cancer originates when a somatic cell deviates from the cooperative regulation of multicellularity and begins to evolve as an independent entity. During cell division, genetic and epigenetic changes occur in the offspring of cancer cells, giving rise to new trait variations and leading to emergence of subclones. Continuous subclonal diversification, also known as intra-tumor heterogeneity (ITH), allows for somatic Darwinian evolution, particularly in circumstances of competition for limited resources among early cancer cells^1–3^. Importantly, the presence of genotypic and phenotypic ITH is associated with poor prognosis and treatment failure ^4,5^. Consequently, determining the fundamental principles governing somatic evolution and the maintenance of ITH in the face of subclonal competition is crucial for developing more effective strategies for tumor prevention and therapy^6^.

The APOBEC (Apolipoprotein B mRNA-editing enzyme catalytic polypeptide-like) genes are a family of enzymes involved in single stranded DNA (ssDNA), and to a lesser extent RNA, deamination of C sites^7–13^. Several members of the APOBEC3 sub-family, particularly APOBEC3A and APOBEC3B are associated with cancer initiation or progression^14^. Many subsequent studies have established APOBEC3A and B as causal for mutations in genomic DNA by targeting ssDNA during the process of replication or transcription ^15,16^. APOBEC3A and B are the only two APOBEC3s that enter the nucleus ^17–19^, and recent work suggests that APOBEC3A may be more mutagenic despite lower expression levels ^20^. APOBEC cytidine deamination activity results in C-to-U conversion and, following replication can generate C-to-T mutations or, more rarely, C-to-A and C-to-G mutations that depend on downstream error-prone DNA repair ^21,22^. These mutations can contribute to the initiation and progression of cancer by disrupting the normal function of genes ^23–25^.

Intra-tumor heterogeneity can provide a basis for the high evolutionary capacity (evolvability) of cancer by increasing the probability of having progeny that can survive substantial selection ^26–28^. In a tumor, there are two types of somatic aberrations: founder mutations and progressor mutations. Founder mutations, also called trunk mutations or clonal mutations, are present in all tumor cells and are caused by an initial carcinogenic event, driving tumor initiation and growth. Progressor mutations, also known as branch/leaf mutations or subclonal mutations, occur later in tumor development, contributing to tumor progression and heterogeneity by introducing additional genetic changes. Distinguishing between these two types of aberrations is vital for understanding the complex genetic landscape and evolution of tumors. Therefore, it is crucial to understand how different mutational processes can generate intra-tumor heterogeneity influencing tumor evolution trajectory^29^.

Analysis of large datasets of genome-level mutations have revealed that diverse mutational processes generate distinct patterns of mutations known as mutational signatures ^30–32^. Alexandrov et al. (2013 and 2020)^30,32^ conducted a comprehensive study on the patterns of somatic mutations in cancer genomes using whole exome sequencing data. Through their analysis, they identified more than 30 distinct mutational signatures, specifically single-base substitution (SBS) signatures. These SBS signatures represent characteristic patterns of DNA base substitutions that occur during the development of different cancers. Each signature is defined by the specific types of base substitutions it comprises and the sequence context in which these substitutions tend to occur. These signatures provide insights into the underlying mutational processes that contribute to cancer development and progression. The mutational signatures encompass a range of mutational processes, including those associated with aging, DNA repair deficiencies, and exposure to various mutagens such as tobacco smoke and ultraviolet radiation. While the etiology of many mutational signatures remains unknown, the causes of many mutational signatures are supported by experiments, including two APOBEC signatures. SBS signatures 2 and 13, which are associated with APOBEC3 induced mutagenesis, have been observed in various types of cancers, including breast, bladder, and lung cancers ^33,34^. The association of these signatures with APOBEC activity has been well-documented in some cancer studies, particularly in early stages of breast and ovarian cancer^35–37^.

We previously developed a program, Cytidine Deaminase Under-representation Reporter (CDUR), that uses computational statistical methods to evaluate previous evolutionary influence of APOBEC on coding sequences of viral genomes ^38,39^. Different APOBEC proteins prefer a cytidine target in different targeting motifs such as T<u>C</u>, TC<u>C</u>, or CC<u>C</u> in case of APOBEC3A/B, 3C, and 3G in single-stranded DNA. The program evaluates whether a coding sequence contains a statistically significant under-representation (scarcity) or over-representation (abundance) of cytidine deaminase mutation motifs. This statistical approach involves creating a null distribution to describe the expected occurrence of mutation motifs within the analyzed sequence (the “subject”) by using codon shuffling methods, for example, rearranging nucleotides at the 3rd codon position, while preserving the amino acid sequence. The subject sequence is then compared to this expected null distribution, and a P value can be calculated for under- or over-representation. In addition to assessing the count of cytidine deaminase mutation motifs, CDUR also computes a related statistic for the number of nonsynonymous mutations occurring at these motifs and the ratio of nonsynonymous mutations to mutation motifs, a measure we describe as mutational susceptibility.

We applied CDUR to study human genes and their relationship to APOBEC motif preferences. In this study, we analyzed motif under-representation and mutational susceptibility of APOBEC3A/B TC hotspots motif of the entire human transcriptome (over 40,000 protein coding transcripts), and found a distribution that is skewed towards APOBEC3A/B robustness, i.e., tolerance of mutations. Furthermore, we selected cancer-associated genes and non-cancer associated (control) genes with no point mutations reported in the COSMIC database. We then compared the distributions of APOBEC3A/B motifs statistics of the two groups and found the two distributions to be very different. The cancer genes had a bimodal distribution containing both under- and over-representation of APOBEC3A/B targeting motifs. We further investigated the APOBEC3A/B motif statistics of orthologs of cancer-associated and control genes in other species. Interestingly, orthologs which, have similar protein sequences and are likely to have the same function, showed a very broad range of motif under-representation and mutational susceptibility, suggesting that genes are labile in terms of these statistics. Our analyses of both human and bat genomes suggested that APOBEC targeting preferences predominantly evolved to avoid excessive damage to the genome, rather than the genome itself. In addition, our simulations suggest that additional APOBEC activity together with the bimodal distribution of susceptibility to APOBEC can introduce higher heterogeneity during the clonal evolutionary process in cancer development.

## Results

### APOBEC3A/B TC hotspots analysis of human genome protein-coding transcripts shows a biased distribution

We hypothesized that APOBEC motif preferences have co-evolved with the human genome. This co-evolution may have caused human genes to evolve so as to avoid APOBEC mutagenesis. At the same time, APOBEC preferences may also have evolved to minimize damage to the human genome. To evaluate the extent to which the human genome will be affected by APOBEC activity, we analyzed how much each gene has significantly fewer mutational motifs (motif under-representation) and depletion of motifs in synonymous positions (defined as the fraction of nonsynonymous mutation motifs, a measure we will refer to as mutational susceptibility) using CDUR ^38^. Figure 1a shows both measures for APOBEC3A/B T<u>C</u> hotspots, for each gene in the human transcriptome (GENCODE v40). Both the motif under-representation and mutational susceptibility measures are heavily skewed, primarily towards the top-left, and to some extent also towards the bottom-right corner of the CDUR plot. A possible explanation for the high density of transcripts in the top-left corner is that a significant number of transcripts have evolved to avoid APOBEC3A/B mutagenesis such that T<u>C</u> motifs have become under-represented in the sequence and those that remain are mostly inevitable at nonsynonymous sites. At the same time the APOBEC3A/B TC motif may have evolved to cause less damage to our genome, especially to crucial genes (e.g., DNA repair). We suggest three possible scenarios to explain the high density in the bottom-right corner: (i) the simplest scenario is that these genes are not exposed to APOBEC3A/B so the T<u>C</u> motif abundance of the genes was not subject to selection by APOBEC3A/B activity. (ii) genes might have followed a distinct strategy to avoid function loss by APOBEC3A/B mutation by increasing the number of TC hotspots and thus making the hotspots less susceptible. (iii) Another scenario is that selection independent of APOBEC3A/B-induced mutagenesis would have led to increased TC hotspots with the underlying cause of decreased mutational susceptibility remaining unknown.

**Figure 1.**
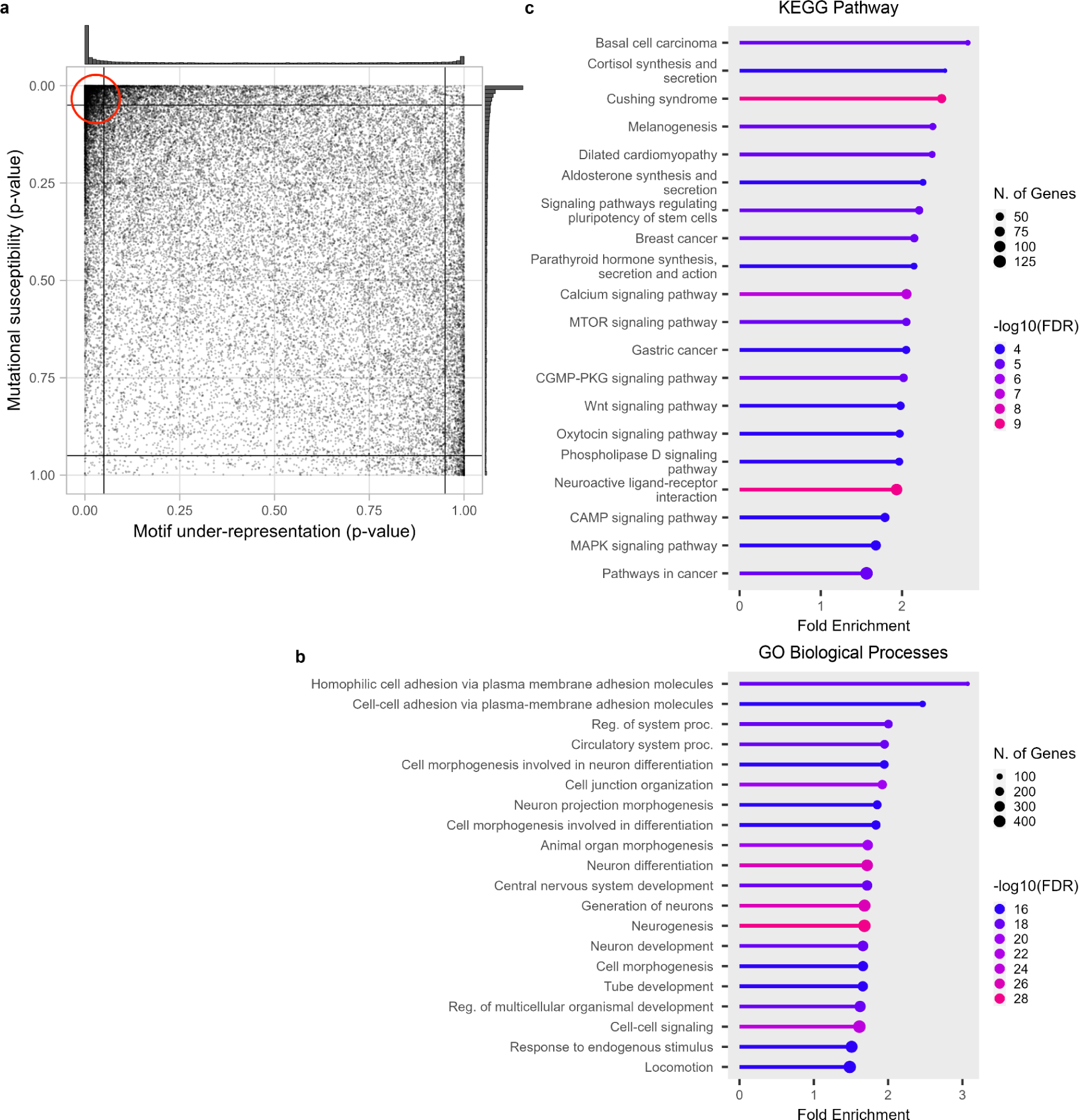
APOBEC3A/B TC hotspots statistics of protein-coding transcripts of the human genome. a) CDUR analysis on human genome shows two dense regions at top-left and right-bottom extremes and relatively sparse dispersion elsewhere. b) Genes in the top-left partition of the CDUR plot showed significant enrichment of GO Biological Processes associated with differentiation and development. c) Genes in the top-left partition of the CDUR plot showed significant enrichment of KEGG Pathway associated with several cancers, basal cell carcinoma, breast cancer and gastric cancer.

### Genes located in top-left corner of the CDUR plot display a significant enrichment of KEGG pathways associated with cancer

To understand the two extreme groups of genes discovered in the CDUR analysis, we performed gene enrichment analysis. First, the top-left corner has 6,539 transcripts of 3,869 genes. Interestingly, Gene Ontology (GO) Biological Processes analysis of these genes showed that 5 out of top 20 ranked by fold enrichment were associated with differentiation and development (cell morphogenesis involved in neuron differentiation, cell morphogenesis involved in differentiation, neuron development, tube development, regulation of multicellular organismal development, epithelium development). These processes are associated with cancer through disruption of differentiation and development-associated genes leading to cancer initiation or at least stimulating cancer progression ^40–42^ (Figure 1b). Interestingly, the enrichment analysis on the KEGG pathway database showed significant enrichment in several cancers (Figure 1c). To confirm the top-left genes association with cancer pathways, we randomly selected the same number of genes outside of the top-left corner 20 times and performed KEGG pathway enrichment analysis. We strictly constrained cancer related pathway as pathways in 6.1 Cancer: overview and 6.2 Cancer: specific types from KEGG pathway Database. With this criterion, the top-left corner has 4 cancer related KEGG pathways and average fold enrichment of 2.14. In contrast, KEGG pathway analysis of randomly selected genes showed 1.05 cancer related KEGG pathways and 11 out of 20 cases showed no cancer related pathway. Moreover, the average fold enrichment of cancer related pathways was 1.46 lower than 1.56, the minimum cancer related pathway fold enrichment in the top left corner. In addition, if we randomly select a square box size of 0.05 in the plot and run KEGG pathway analysis, it did not show any significant fold enrichment. (Figure S10). For GO Cellular Components, enrichment analysis showed a significant fold enrichment of channel complex and synapse components (Figure S1). Enrichment analysis on GO Molecular Function term showed a significant fold enrichment of mostly ion channel or transporter activity. This is an intriguing result considering the role of ion channels and transporters in solid tumors, and their adaptation to harsh microenvironments such as hypoxia and acidosis^43^. Second, the bottom-right corner has 396 transcripts of 230 genes (Figure S2). GO Biological Processes on these genes did not show any significant enrichment. KEGG pathway enrichment showed herpes simplex virus 1 (HSV-1) infection (Figure S3). HSV-1 genome is potential substrate of APOBEC3 so abundant TC hotspots with low susceptibility of those genes would be expected. Gene enrichment analysis on GO Cellular Component showed mostly collagen-associated terms (Figure S4), and GO Molecular Functions showed mostly DNA binding activity (Figure S5).

### Cancer-associated genes have a distinct bimodal distribution of APOBEC3A/B under-representation

As the KEGG pathway enrichment analysis of the genes in the top-left corner showed a significant fold enrichment of cancer-associated pathways, we further analyzed known cancer-associated genes. We used the COSMIC database list of cancer-associated genes, which were retrieved from the COSMIC Cancer Gene Census dataset ^44,45^. The COSMIC Cancer Gene Census are cancer-associated genes systematically curated by professional researchers. For comparative purposes, we generated a list of control genes collecting protein-coding genes from the NCBI Gene database that do not have any annotated point mutations in the COSMIC Mutation database. While applying the method, we observed that the number of control genes was significantly lower than that of cancer-associated genes. Nevertheless, to maintain a systematic analysis, we retained this criterion. Figure 2a shows the same data as Figure 1a, but with the cancer genes highlighted in red and the control genes in blue. Here we found a significant difference in the distribution between cancer-associated genes and the control genes group in the CDUR plot (Figure 2a) (Kolmogorov-Smirnov test for motif-representation axis, P = 2.828 × 10^-7^; for mutational susceptibility axis, P = 1.175 × 10^-4^). However, in the motif under-representation axis, a difference of means test did not show a significant difference between the two groups (Wilcoxon rank sum test, P = 0.1047) due to the bimodal form of the cancer gene distribution compared to the unimodal control gene distribution. Along the mutational susceptibility axis, cancer-associated genes do have a significantly higher mutational susceptibility than control group genes (P = 4.838 × 10^-5^). In summary, cancer associated genes have a bimodal distribution for TC motif under-representation that is distinct from the control genes. Cancer associated genes also have significantly higher mutational susceptibility (y-axis of Figure 2a), consistent with a depletion of TC hotspots in synonymous sites. We compared genes from sex chromosomes and found no significant difference between × and Y chromosomes (Kolmogorov-Smirnov test for motif-representation axis, P = 0.1838; for mutational susceptibility axis, P = 0.4523, Wilcoxon rank sum test for motif-representation axis, P = 5.515 × 10^-2^; for mutational susceptibility axis, P = 0.6658). However, the difference between them is inconclusive since the number of genes between × (1,676) and Y (83) chromosome is large (Figure S8).

**Figure 2.**
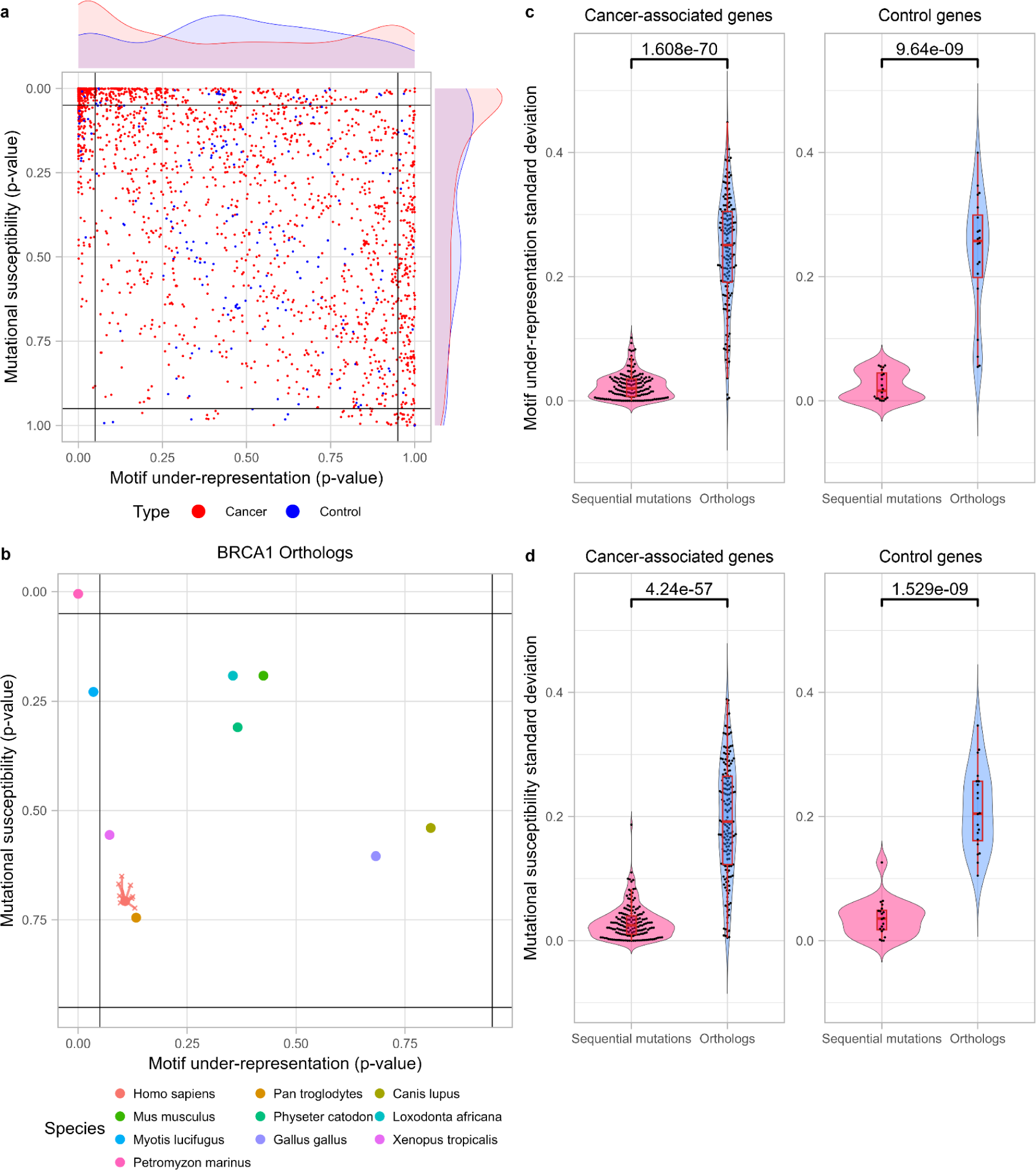
Cancer-associated genes have significantly different distribution in CDUR plot compared to control genes. a) Cancer-associated genes and control genes show distinct distribution in both motif under-representation and mutational susceptibility (Kolmogorov-Smirnov test for motif-representation axis, P = 2.828 × 10^-7^; for mutational susceptibility axis, P = 1.175 × 10^-4^). b) Motif under-representation and mutational susceptibility of BRCA1 orthologs show higher variance than sequential mutations of the gene on CDUR plot. c) Standard deviation of motif under-representation comparison between orthologs and sequential mutations show significantly high variance among orthologs in both cancer-associated genes (196 genes) and control genes (20 genes). d) Standard deviation of mutational susceptibility comparison between orthologs and sequential mutations show significantly high variance among orthologs in both cancer-associated genes (196 genes) and control genes (20 genes).

### CDUR statistics of orthologous genes from other species are dispersed over a broad range

Given the surprising result of a bimodal distribution for under-representation for cancer associated genes, we sought to investigate the extent of natural variation that is possible in these genes. For this we used the same CDUR measures to consider orthologous genes (orthologs). Orthologs are homologous genes in different species that have highly similar protein sequences. Barring rare cases of convergent evolution, orthologs are derived from a common ancestor gene and have likely preserved the same function. Orthologs are therefore good candidates to evaluate the genotypic heterogeneity of a phenotype. We collected orthologs of 196 cancer-associated genes and 20 control genes from ten vertebrate species including human (*Homo sapiens*, *Pan troglodytes*, *Canis lupus familiaris*, *Gallus gallus*, *Loxodonta africana*, *Mus musculus*, *Physeter catodon*, *Xenopus tropicalis*, *Myotis lucifugus*, and *Petromyzon marinus*) and calculated the CDUR statistics as we did for the human genes. As an example, Figure 2b shows ten orthologs of breast cancer type 1 susceptibility protein (BRCA1), a well-known tumor suppressor gene that is crucial to DNA damage repair, which we found were dispersed across a wide range in the CDUR plot. To investigate the generality of this observation, we compared variation in the CDUR plots of each ortholog set to sequences generated from the corresponding human genes by sequentially mutating T<u>C</u> motifs until we encounter an amino acid change, thus simulating the variation that might be introduced to the human gene by APOBEC3A/B. We found that, regardless of whether the genes are associated with cancer or not, most orthologs showed a much greater variation for both TC motif under-representation and mutational susceptibility when compared to the variation that might be introduced by APOBEC to the human gene (Figure 2c, 2d). Although most of the observed variance in the orthologs is likely unrelated to co-evolution with APOBEC (since there are many other evolutionary pressures involved) the broad range distribution suggests that genes at least have the potential to evolve a wide range of tolerance to APOBEC3A/B-induced mutagenesis while maintaining their essential functions.

### TC hotspot analysis of the bat genome suggests predominant evolution of APOBEC preferences rather than genomes

To further investigate the relationship between distribution of TC hotspots in the human genome and APOBEC3A/B motif preference, we performed the same TC hotspots analysis of a bat (*Pteropus alecto*) genome. *Pteropus alecto* has undergone an expansion of APOBEC3 genes, leading to 18 APOBEC3 genes, compared to 7 in humans. As in humans, the targeting preference is also predominantly T<u>C</u> ^46^. Bats are known to be long-lived animals and have low cancer incidence^47,48^. If APOBEC were a main cancer initiating factor in bats, we would expect to observe a different TC hotspot distribution compared to the human genome. Interestingly, the *Pteropus alecto* genome displayed a similar overall pattern to the human genome; high density in top-left and bottom-right corner (Figure 3a; Figure S7). Despite the qualitative similarity in both axes between the human and bat genome distributions, there is a greater under-representation of TC hotspots in the human genome (Willcoxon rank sum test for motif under-representation: P < 2.2 × 10^-16^ and for mutational susceptibility: P = 2.204 × 10^-10^). We also analyzed the distribution of bat orthologs of both human cancer-associated genes and control genes (Figure 3b), and again observed similar features to the human distribution, including the bimodal distribution for TC hotspot under-representation. Furthermore, we analyzed 5 different species including elephant, chimpanzee, mouse, yeast, and nematode worm (Figure S9). We found no qualitative difference between mammalian species. The yeast genome showed a uniform distribution in both measurements. The C. elegans genome showed that most genes have over-representation of TC hot spots and low mutational susceptibility. We are interested in bats because of their longevity despite small size and because the APOBEC gene family has expanded in various species. Compared to bats, all other species studied have fewer APOBECs, yet results from bats were not qualitatively different than those in other species. These results suggest that the APOBEC TC motif preference may have evolved to avoid harm to its genome, but the genome itself does not appear to have been greatly shaped by APOBEC activity.

**Figure 3.**
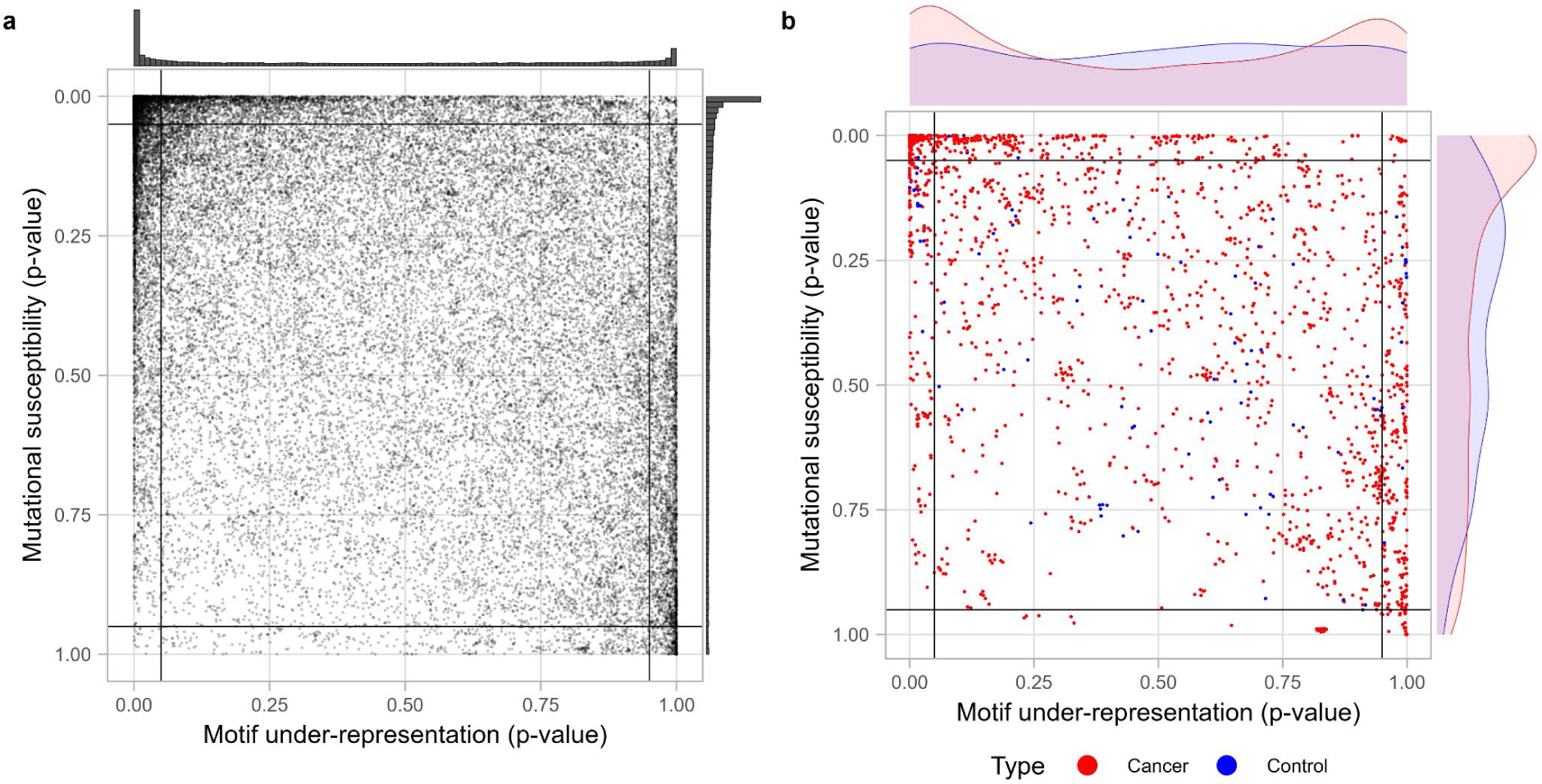
APOBEC3A/B TC hotspots statistics of protein-coding transcripts of *Pteropus alecto* genome. a) CDUR analysis on bat genome shows two dense regions at top-left and right-bottom extremes and relatively sparse dispersion elsewhere as in the human genome. b) Bat orthologs of human cancer-associated genes and control genes show distinct distribution in mutational susceptibility but not in motif under-representation and (Kolmogorov-Smirnov test for motif-representation axis, P = 0.1701; for mutational susceptibility axis, P = 3.169 × 10^-5^)

### Role of APOBEC3A/B in intra-tumor heterogeneity

In order to better understand the consequence of the bimodal distribution of APOBEC3A/B motif under-representation on tumor evolution and heterogeneity, we performed simulations. We adapted a previous computational model constructed with the Java based spatial platform HAL ^49,50^ to run simulations comparing clonal dynamics and trajectories over tumor evolution. Briefly, the model starts with cells without any mutations and at each timestep cells can divide to produce additional single cells or die. Depending on the average ratio between division rate and death rate in a population, the number of cells in a population may increase, decrease, or stay the same. A cell can acquire a single mutation according to the given mutation rate when the cell divides. The mutation can affect the fitness of cells positively, neutrally, and negatively, and the division rate at the next time step will be updated accordingly. First, we compared simulations with only random mutations and with additional APOBEC3A/B mutations where all genes have the same probability to get positive and negative mutations, to evaluate if APOBEC activity alone can increase heterogeneity (Figure 4a and 4b). We found that although APOBEC3A/B activity increases overall mutation rate, there is no significant difference in intra-tumor heterogeneity (Figure S6a and S6b). Muller plots from each condition show that intra-tumor heterogeneity increases faster when there is APOBEC3A/B, however the intra-tumor heterogeneity difference at the final time point is negligible (Figure 4c and 4d). Then we simulated the effect of bimodal distribution of APOBEC3A/B motif under-representation that was shown in Figure 1a. We showed above that the bimodal distribution is more evident in cancer-associated genes compared to control genes. To investigate the role of the bimodal distribution, we assumed three categories of genes based on their motif under-representation. The genes with “low” and “high” motif under-representation are defined as genes whose motif under-representation is below 0.05, or above 0.95, respectively, and those that fall in between are defined “mid”. Then, we assigned the probability of genes to fall into each category according to the distribution of interest. For example, if we assume motif under-representation of a genome is uniformly distributed, the probability that a gene falls into the “low” or “high” category will be equal to its proportion, 0.05, and 0.9 for the “mid” category. In contrast, if we assume a bimodal distribution based on Figure 1a, the probability for “low”, “mid”, and “high” becomes 0.3, 0.6, and 0.1, respectively. In addition, we set a higher probability to get a positive mutation than negative mutation for the genes in the “low” and “high” category to emulate the result of Figure 2a that shows cancer-associated genes mostly fall into “low” and “high” category. During the simulation we recorded the number of genotypes in the population and computed heterogeneity based on Shannon entropy as in West et al. 2021^50^. We found the bimodal distribution leads to higher intra-tumor heterogeneity (Figure 4e and 4f). Furthermore, the difference of means test showed increasing -log(P) value indicating that the difference became substantial (Figure S6c and S6d). We found no evidence of linear or branching evolution in our simulation, and we only observed neutral evolution over 600 time steps. Based on our simulations, the effect of APOBEC activity on tumor initiation is comparable to random mutation, however, APOBEC activity with bimodal distribution of TC hotspots in genome can increase final tumor heterogeneity.

**Figure 4.**
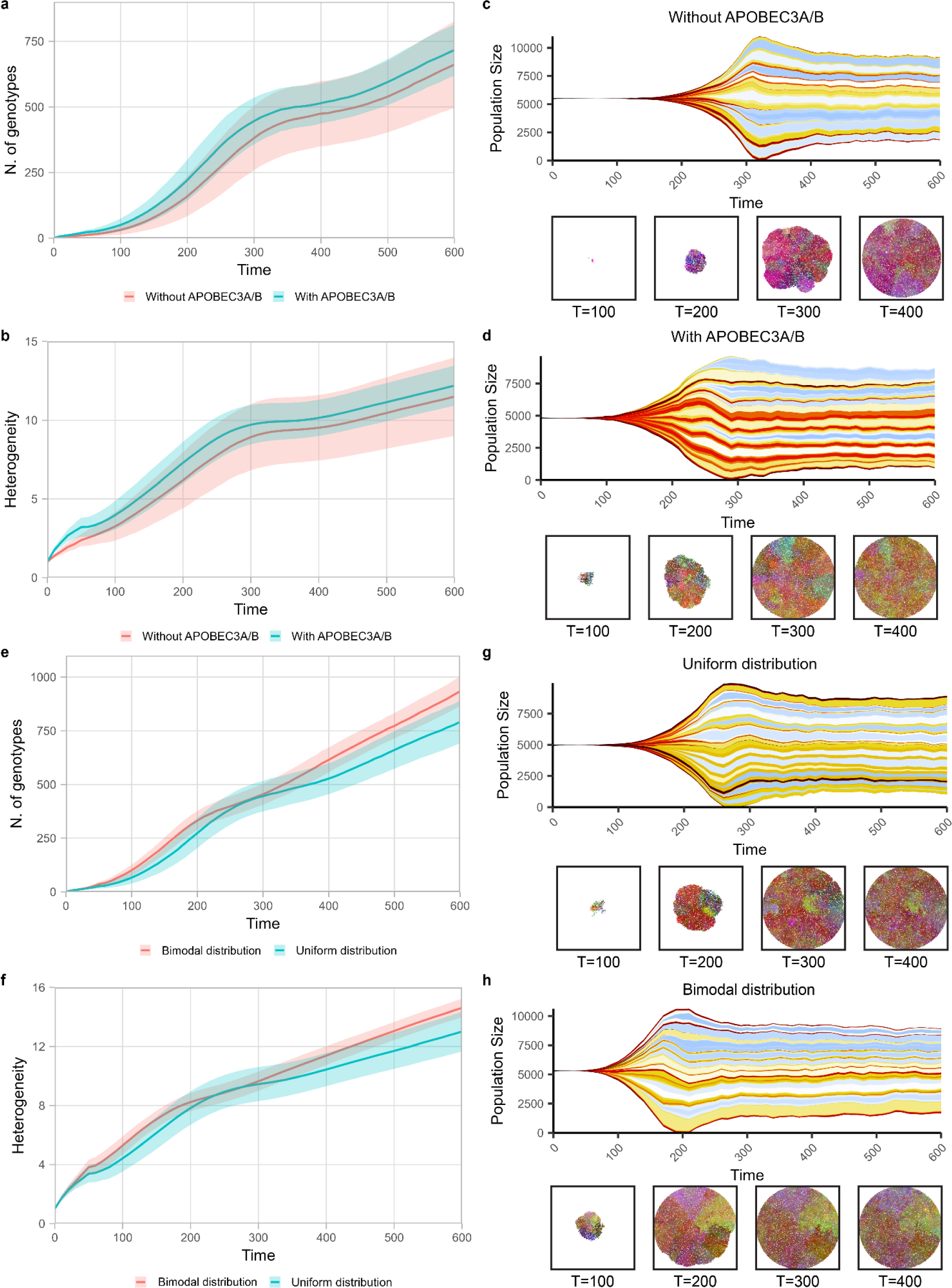
Protein-coding transcripts from human genome display biased distribution in APOBEC3A/B motif representation and mutational susceptibility. Bimodal distribution of APOBEC3A/B motif representation displays higher heterogeneity by spatial clonal dynamics simulation. a) The number of genotypes between with and without APOBEC3A/B shows no significant difference. b) Heterogeneity between the two simulations with and without APOBEC3A/B shows no significant difference. c, d) Muller plots of simulation with and without APOBEC3A/B activity shows no significant difference in heterogeneity at the final time point. e) The number of genotypes between uniform and bimodal distribution shows an increasing difference over time. f) Heterogeneity between uniform and bimodal distribution shows an increasing difference over time. g, h) Muller plots of simulation uniform and bimodal distribution show an increasing difference over time.

## Discussion

Evolvability is an emerging hallmark of cancer. The presence of genetic and phenotypic intratumor heterogeneity (ITH) within a tumor is the main source of evolvability that has also been recognized as a significant factor contributing to poor prognosis and treatment failures in cancer patients. Several studies have highlighted the impact of ITH on disease progression and clinical outcomes ^4,5^. The heterogeneity itself arises through ongoing somatic evolution, driven by genetic mutations and selection on clones. Different subclones within the tumor can possess distinct genetic alterations represented by a selectable phenotype, leading to variability in their response to their variable tumor microenvironment and increasing tumor fitness in general. Consequently, some subclones may survive and become dominant, while others may be resistant or susceptible to specific selection by tumor microenvironments. Understanding the fundamental principles that govern somatic evolution and the maintenance of ITH is crucial to understand tumor initiation, progression, evolution, and evolvability. However, the main question that is still unanswered is how tumor heterogeneity fuels tumor evolution leading to increased cancer cell evolvability.

To answer that question, we designed an analysis pipeline for measuring genome mutation tolerance considering APOBEC as a mutator. The idea was how random mutation can affect genotypic heterogeneity while maintaining a protein-coding phenotype. We used CDUR to analyze the human genome with respect to APOBEC mutation potential. We analyzed motif under-representation and mutational susceptibility of APOBEC3A/B TC hotspots motif of more than 40,000 human protein-coding transcripts. CDUR analyses revealed an evident disproportion in both motif under-representation and mutational susceptibility. Transcripts of the human genome were densely concentrated in the top-left or bottom-right corner of the plot, indicating distinct strategies employed by genes to either avoid APOBEC3A/B mutagenesis or increase the number of TC hotspots while reducing mutational susceptibility ^39,51^. However, it is also likely that APOBEC evolved to avoid causing damage to genes rather than genes evolving to avoid the hotspots. Despite having many more APOBEC3 genes, hotspot under-representation in *Pteropus alecto* mirrored that of humans. Similarity in pattern suggests APOBEC evolution to mitigate impacts on the genome instead of the converse.

Moreover, genes located in the top-left corner of the CDUR plot exhibited significant enrichment in cancer-associated pathways, suggesting a potential link between APOBEC3A/B TC hotspots statistics and cancer formation. However, it is still not quite clear the extent to which human genes may have evolved under APOBEC selection and how this adaptation to APOBEC is related to their original role in tissue or tumors or correlated with the tumor microenvironment.

Further analysis of cancer-associated genes and control genes group on the CDUR plot demonstrated a distinct distribution compared to control genes. Cancer-associated genes exhibited significantly higher mutational susceptibility. The presence of two peaks at opposite positions in the motif under-representation axis likely contributed to the lack of significant differences in mean between the two groups. However, the Kolmogorov-Smirnov test confirmed that cancer-associated genes and control group genes differed significantly in both axes of the CDUR plot.

Additionally, the analysis of orthologs from multiple vertebrate species revealed a wide range of variance in motif representation and mutational susceptibility for genes, irrespective of their association with cancer. This observation suggests that genes undergo various evolutionary trajectories and can potentially greatly alter the representation and susceptibility of APOBEC3A/B motifs. The broad range distribution of orthologs in the CDUR plot indicates that there can be multiple strategies genes might employ to tolerate APOBEC3A/B-induced mutagenesis and maintain their essential functions. Our simulations showed the effects of APOBEC-driven heterogeneity on cancer cell clonal dynamics and evolutionary trajectories. We found APOBEC can significantly influence cancer-associated genes compared to control genes, leading to a notable increase in intra-tumoral heterogeneity in a neutral evolution manner. Neutral evolution refers to the maintenance of genetic changes that arise randomly without conferring a selective advantage or disadvantage to cells. In cancer, substitution accumulation can occur in two ways: some mutations provide a selective advantage, leading to clonal expansion and dominant subclones, while others arise randomly and have minimal impact on cell fitness. Over time, and with stable or increasing cell population size, these neutral substitutions accumulate, contributing to genetic diversity and intra-tumor heterogeneity. In the context of APOBEC activity, we propose two effects on evolution. Firstly, APOBEC activity increases the occurrence of random mutations within the genome. These substitutions result from the enzymatic activity of APOBEC. Secondly, APOBEC activity exhibits a biased increase in mutations within a subset of cancer-related genes compared to non-cancer genes. This bias towards cancer-associated genes further shapes the genetic landscape of the tumor, potentially influencing tumor progression and response to treatment. In summary, APOBEC activity impacts neutral evolution through increased random mutations that generates greater variation, and a preferential increase in mutations within cancer-related genes. Both effects contribute to the genetic diversity and intra-tumor heterogeneity observed in cancer.

Overall, our study provides insights into the APOBEC3A/B TC hotspots in the human genome and highlights the distinct characteristics of cancer-associated genes in the APOBEC3A/B motifs statistics.

These findings contribute to our understanding of the interplay between APOBEC3A/B mutagenesis and the development of cancer. We found APOBEC has a significantly stronger influence on progressor genes and mainly on cancer-associated genes compared to control genes, leading to increased intra-tumoral heterogeneity. Our findings emphasize the critical role of APOBEC as an accelerator rather than initiator in carcinogenesis, which further amplifies intra-tumoral heterogeneity toward neutral evolution.

### Limitations of study

This study has several limitations to consider. Firstly, it focuses solely on APOBEC3A/B TC hotspots and does not account for other mutational processes or factors that may contribute to the observed patterns. Secondly, the reliance on computational analyses and statistical tests introduces inherent assumptions and potential inaccuracies. Furthermore, the study’s conclusions are based on correlations rather than causation, and experimental validation is necessary. The use of CDUR statistics as the primary analysis tool has its own limitations, and alternative or complementary approaches could provide additional insights. Lastly, while the comparative analysis of orthologs across species is informative, species-specific factors may influence mutational dynamics. Overall, while valuable, the findings should be interpreted cautiously, and further research is needed to address these limitations and provide a more comprehensive understanding of the relationship between mutational processes, gene function, and cancer.

## STAR ★ Methods

### Key resources table

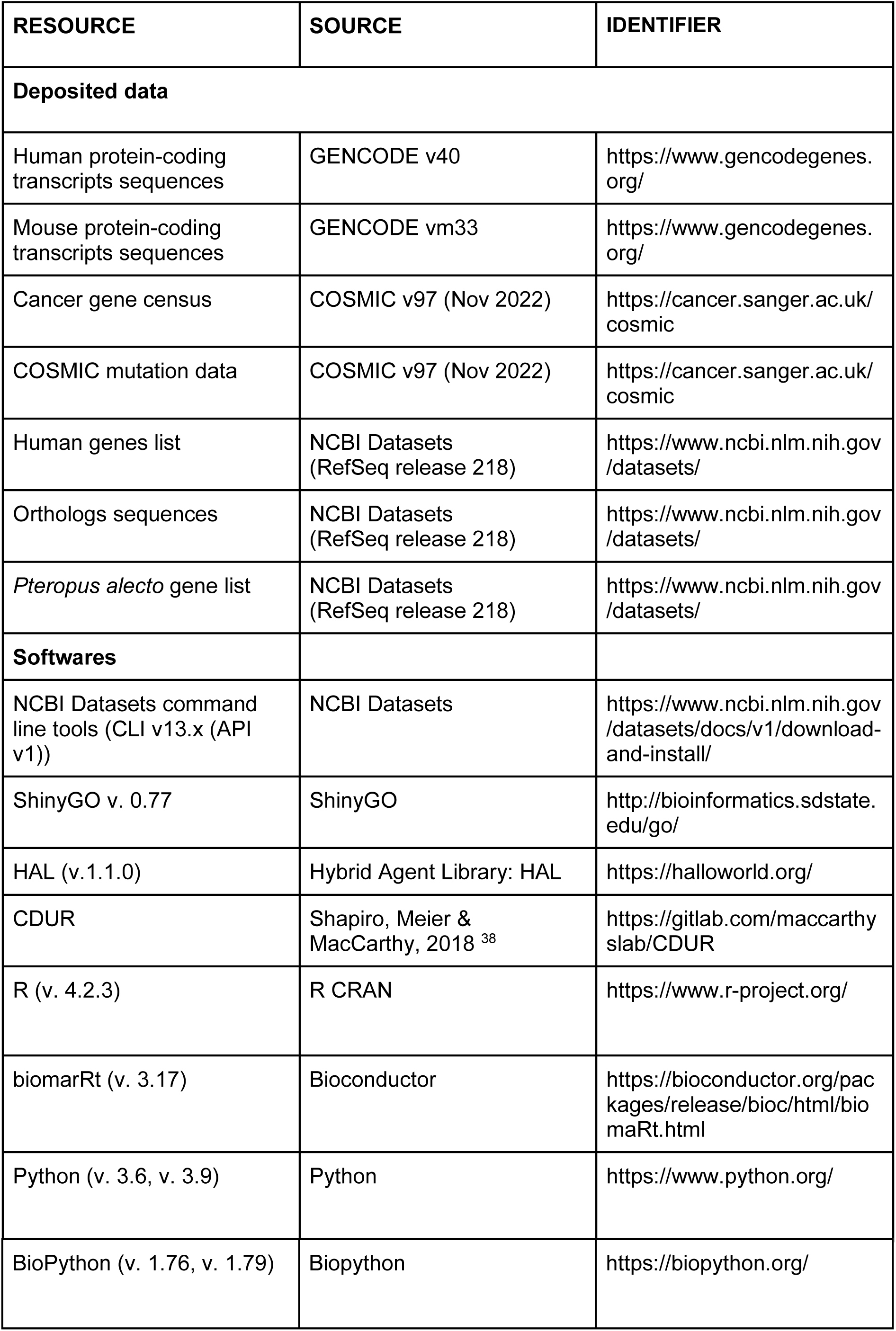

## Resource availability

### Lead contact

Further information and any related requests should be directed to and will be fulfilled by the lead contact Mehdi Damaghi and Thomas MacCarthy.

### Materials availability

This study did not generate new unique reagents.

## Method Details

### Transcripts sequences preprocessing

CDUR requires inputs as FASTA file with each record being a complete coding sequence starting with the start codon and ending with stop codons. The FASTA file from GENCODE contains protein coding transcripts sequences that have not only coding sequences but also 5’ and 3’ untranslated regions. In addition, several sequences were incomplete coding sequences. To make analysis only on valid coding sequences, we filtered records in the original GENCODE FASTA file and extracted valid sequences. First, substrings of sequence were extracted based on annotation in FASTA file name field. Then, we checked four criteria; 1) whether the length of coding sequence is multiple of three, 2) whether the sequence starts with ATG, 3) whether the sequence ends with stop codon, and 4) whether the name contain PAR_Y indicating pseudoautosomal region from Y chromosome. Transcript sequences that passed all criteria were saved in single FASTA for CDUR input. Aforementioned preprocessing was applied to human and mouse genome analysis.

For other species including black fruit bat (*Pteropus alecto*), elephant (*Loxodonta africana*), chimpanzee (*Pan troglodytes*), yeast (*Saccharomyces cerevisiae*), and nematode worm (*Caenorhabditis elegans*), we used NCBI Datasets command line tools to download all annotated genes information. Then we used a custom Python script with BioPython Entrez packages to retrieve coding sequences of all valid transcript sequences. The scripts first utilized previously downloaded genes information to retrieve all protein-coding NCBI gene id. In the second step, the script efetchs gene ids to get associated RefSeq ID. Third, RefSeq IDs were fetched to get GenBank information and coding sequences were extracted based on the information. Finally, all valid transcripts sequences were saved in a single FASTA file for CDUR input.

### APOBEC3A/B motifs statistics analysis with CDUR

We utilized the Cytidine Deaminase Under-representation Reporter (CDUR) to evaluate APOBEC3A/B motifs statistics, motif under-representation and mutational susceptibility, of human transcripts. Briefly, CDUR generates a null model by generating shuffled sequences from a given sequence of interest. Then CDUR computes motif under-representation of the fraction of sequence in the null model which has fewer motifs than given sequence. For TC hotspots, we used motif under-representation as “belowT_C_’’ value from CDUR result. Mutational susceptibility was defined by how probable the sequence is for transition mutations in hotspots to result in amino acid alterations. CDUR results contain a value “repTrFrac_belowT_C_” which is the ratio of non-synonymous change in TC hotspots, and we used 1-“repTrFrac_belowT_C_” for mutational susceptibility. Preprocessed coding sequences (CDS) of transcripts for human and bat were passed to CDUR with default settings; the shuffle method is n3 and the number of shuffles is 1,000. We ran CDUR three times on the same sequences, respectively, and the average of the values was used in analysis.

### Gene enrichment analysis

Top-left genes were defined as genes with under-representation ≤ 0.05 and mutational susceptibility ≤ 0.05. Bottom-right genes were defined as genes with under-representation ≥ 0.95 and mutational susceptibility ≥ 0.95. top-left and bottom-right genes were extracted using dplyr R package. Only gene names were saved in separate text file and submitted to ShinyGO 0.77^52–54^ with default parameters; FDR cutoff = 0.05, # pathways to show = 20, Pathway size: Min = 2 and Max = 2,000. We downloaded the enrichment results of the top 20 pathways from GO Biological Process, GO Cellular Component, GO Molecular Function, and KEGG, and bar plots were generated with R package dplyr and ggplot2 using custom R scripts.

### Cancer-associated genes and control genes

We defined cancer-associated genes as genes in the COSMIC Cancer Census^45^. We retrieved RefSeq ID and Ensembl ID with biomaRt to map these genes to GENCODE transcripts. Bat equivalent cancer-associated genes were collected using NCBI Datasets command line tools with Gene ID extracted from COSMIC Cancer Census. Control genes were defined as genes rarely get point mutations in the coding region of their transcript. We used custom R and shell scripts for the process to filter out those genes. First, all human reference genes were downloaded from NCBI using NCBI Datasets Command line tools. Then all genes from the NCBI list were checked whether genes have been recorded in COSMIC mutation data at least once. Among the genes that did not have any record, we filtered only protein coding sequences. In addition, cloned genes such as gene names starting with LOC were removed, and also mitochondrial genes were removed. We applied the same procedure to collect bat equivalent cancer-associated genes to generate bat equivalent control genes.

### Ten species ortholog genes

Orthologs of human cancer-related or control genes were downloaded from NCBI using NCBI Datasets command lines tool. For each human gene, we retrieved gene id, and used the gene ID to download orthologs. This process downloads all orthologs available. After downloading all orthologs, we selected genes that have orthologs for all ten species. The sequences of the orthologs were preprocessed to pass into CDUR as we did for *Pteropus alecto* genome, and CDUR statistics were computed three times per each sequence. For variation analysis, we generated mutated sequences from human transcript sequences by simulating sequential C-to-T mutation until the amino acid sequence changes. We performed ten simulations for each transcript and standard deviation on both axes were computed.

### Heterogeneity comparison simulation

Clonal expansion simulations were performed using a computation model constructed with Java based spatial simulation platform HAL^49^. Basic scheme of our model was adopted from West et al. 2022^50^. We added more parameters and changed the mutation method to capture the effect of human genome distribution by the APOBEC3A/B motif proportion of each gene. Model starts with 100 cells in the center of a circle radius of 100 grid points where each single cell occupies a single grid point. Each cell in our model can have three different types of mutations that have different effects on the fitness (division rate); positive, neutral, and negative mutations. Positive and negative mutations increase or decrease fitness respectively, and neutral mutations do not have any effect on fitness. During each time step, each cell can divide producing one additional cell into a grid or die off introducing a single empty grid point. This process may increase the number of cells in a population, if the average division rate is higher than death rate, and decrease in opposite situation. Probabilities to divide or die were computed using following equation:

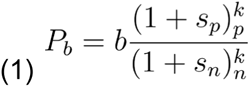

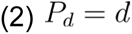

where b and d are the baseline division and death rates, respectively.

The following parameters were used: total number of mutable sites, T_m_=5×10^6^, positive fitness effect, s_p_=0.1, negative fitness effect, s_n_=10^-3^, random mutation rate, μ=10^-8^, APOBEC3 mutation rate μ_a3_=10^-6^. APOBEC3 mutation was turned on from time point 0 to 50 to reflect burst expression in early stage. We used 0.001, 0.01, and 0.1 for the proportion of APOBEC3 for “low”, “mid”, and “high”. To emulate the phenomenon that cancer-associated genes (higher probability to have positive fitness effect) were located in bimodal extreme (“low” and “high”), we set the probability to have mutations that have positive, neutral, and negative fitness to 0.4, 0.5, and 0.1, respectively, and for control genes (higher probability to have negative fitness effect), we set the probabilities to 0.1, 0.5, and 0.4, respectively.

To count the number of genotypes in the population, the genotype ID of the cell at each time point was tagged using the number of each mutation types by equation (3),

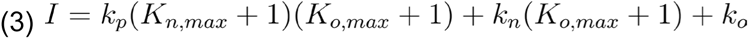

where kp, kn, and ko are the number of positive, negative, and neutral mutations, respectively, and KnMAX and KoMAX are the maximum number of negative and neutral mutations in the population. In addition, heterogeneity were measured using Shannon entropy as in West et al. 2022, given by:

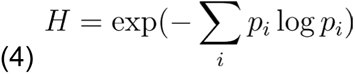

where p_i_ is the proportion of cells within the population with genotype ID is i.

We performed 100 simulations for each condition and used R base package t-test for difference of means test for every 50 time points between with and without APOBEC activity, or uniform and bimodal distribution.

## Supporting information

Suuplementary Figures

## Data and code availability

GENCODE transcripts sequences: All human coding sequences used in this work can be acquired from GENCODE (https://www.gencodegenes.org/).

Cancer mutations data: Cancer related genes and all point mutation data in cancer can be acquired from COSMIC Database (https://cancer.sanger.ac.uk/cosmic).

Orthologs sequences and annotations: Orthlogs RefSeq ID and sequences can be acquired via NCBI Datasets command line tools (https://www.ncbi.nlm.nih.gov/datasets/docs/v2/download-and-install/).

*Pteropus alecto* sequences and annotations: All Pteropus transcripts sequences and annotations can be acquired via NCBI Datasets command line tools (https://www.ncbi.nlm.nih.gov/datasets/docs/v2/download-and-install/).

Clonal evolution simulations of cancer: Hybrid Automata Library (HAL) is freely available from https://halloworld.org/ and the model Java scripts is available in following GitHub repository.

The cytidine deaminase under-representation reporter (CDUR): CDUR is freely available from GitLab repository (https://gitlab.com/maccarthyslab/CDUR).

All original scripts to download and process the data is available at https://github.com/StudyingAnt/Cancer_genes_analysis_of_APOBEC_motifs and are publicly available as of the date of publication.

Any additional information required to reanalyze the data reported in this paper is available from the lead contact upon request.

## Acknowledgements

We gratefully acknowledge funding from Physical Sciences Oncology Network at the National Cancer Institute (grant U01CA261841) as well as support from the Stony Brook Cancer Center. This work was also supported partly by R01 grant R01CA272601. LMD was supported in part by NSF-IOS 2032063.

## Author contributions

J-H. S., T.M., and M.D., designed research; J-H.S performed research; J-H.S contributed new reagents/analytic tools; J-H.S. analyzed data; L.M.D. contributed to results interpretation; and J-H.S., T.M., and M.D. wrote the paper. All authors revised the paper.

## Supplemental information

### Supplementary figures

**Figure S1.**
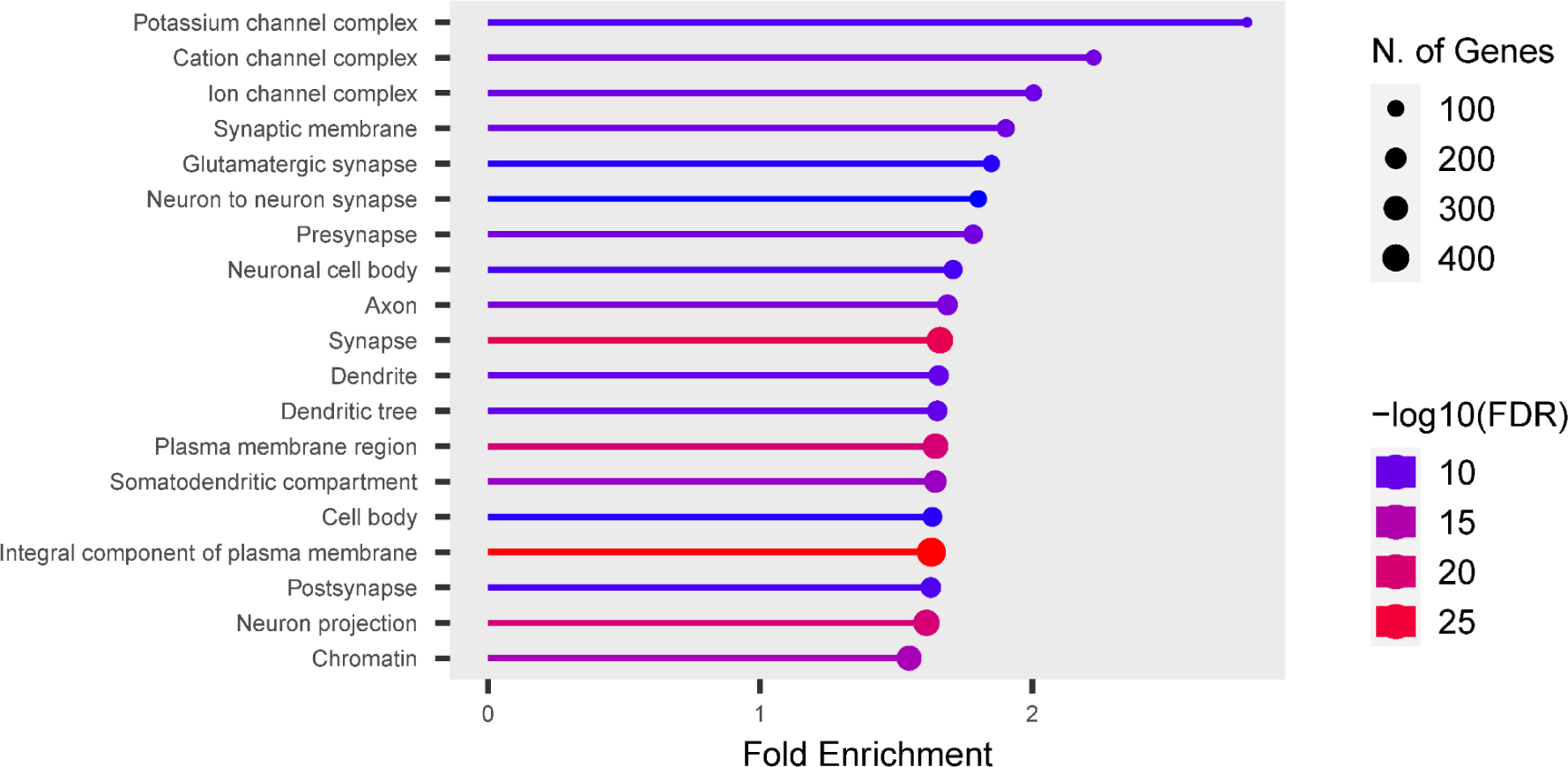
Gene enrichment analysis of GO Cellular Component on top-left corner in human APOBEC3A/B TC motif CDUR plot of gene.

**Figure S2.**
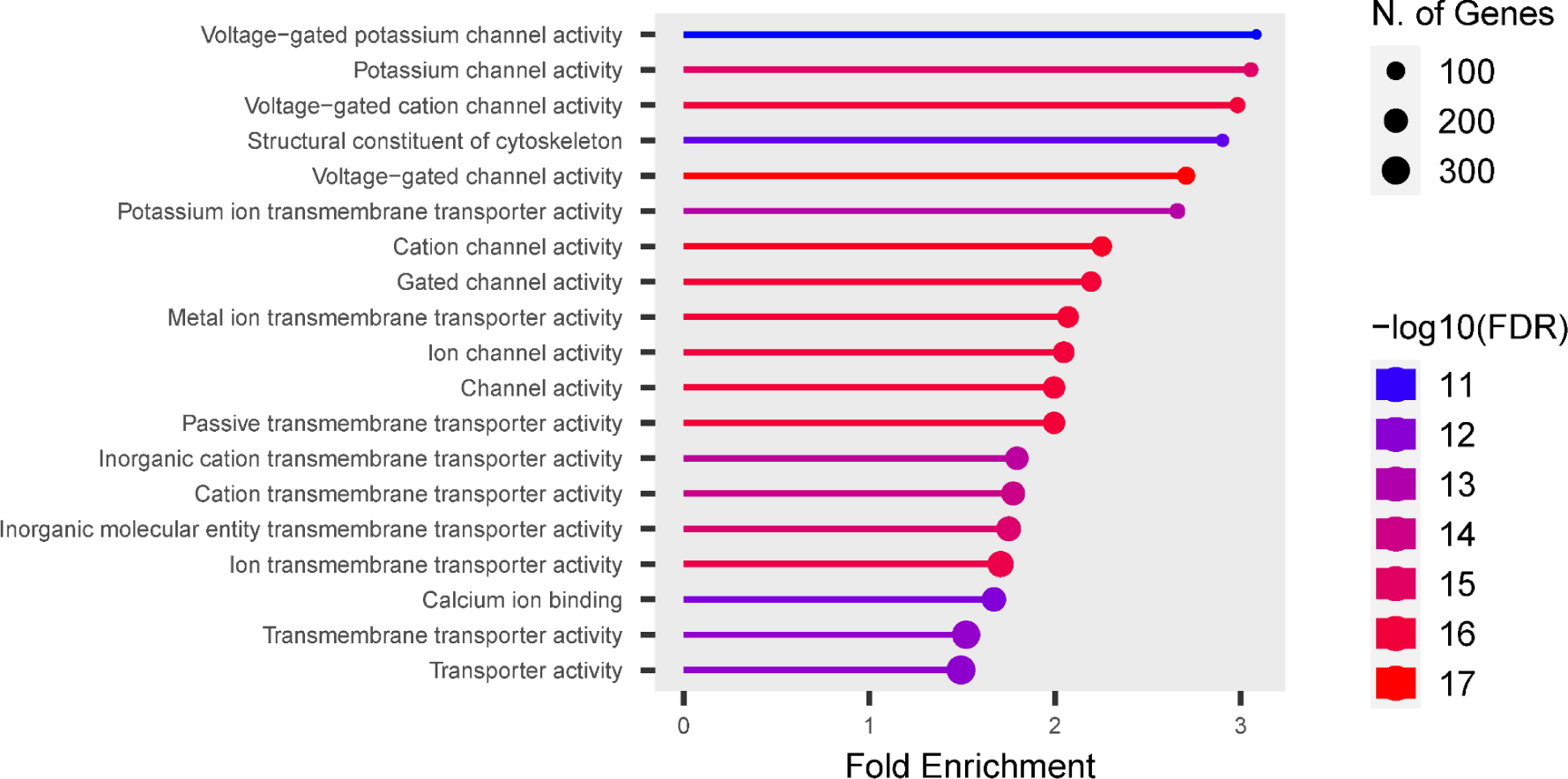
Gene enrichment analysis of GO Molecular Functions on top-left corner in human APOBEC3A/B TC motif CDUR plot of gene.

**Figure S3.**
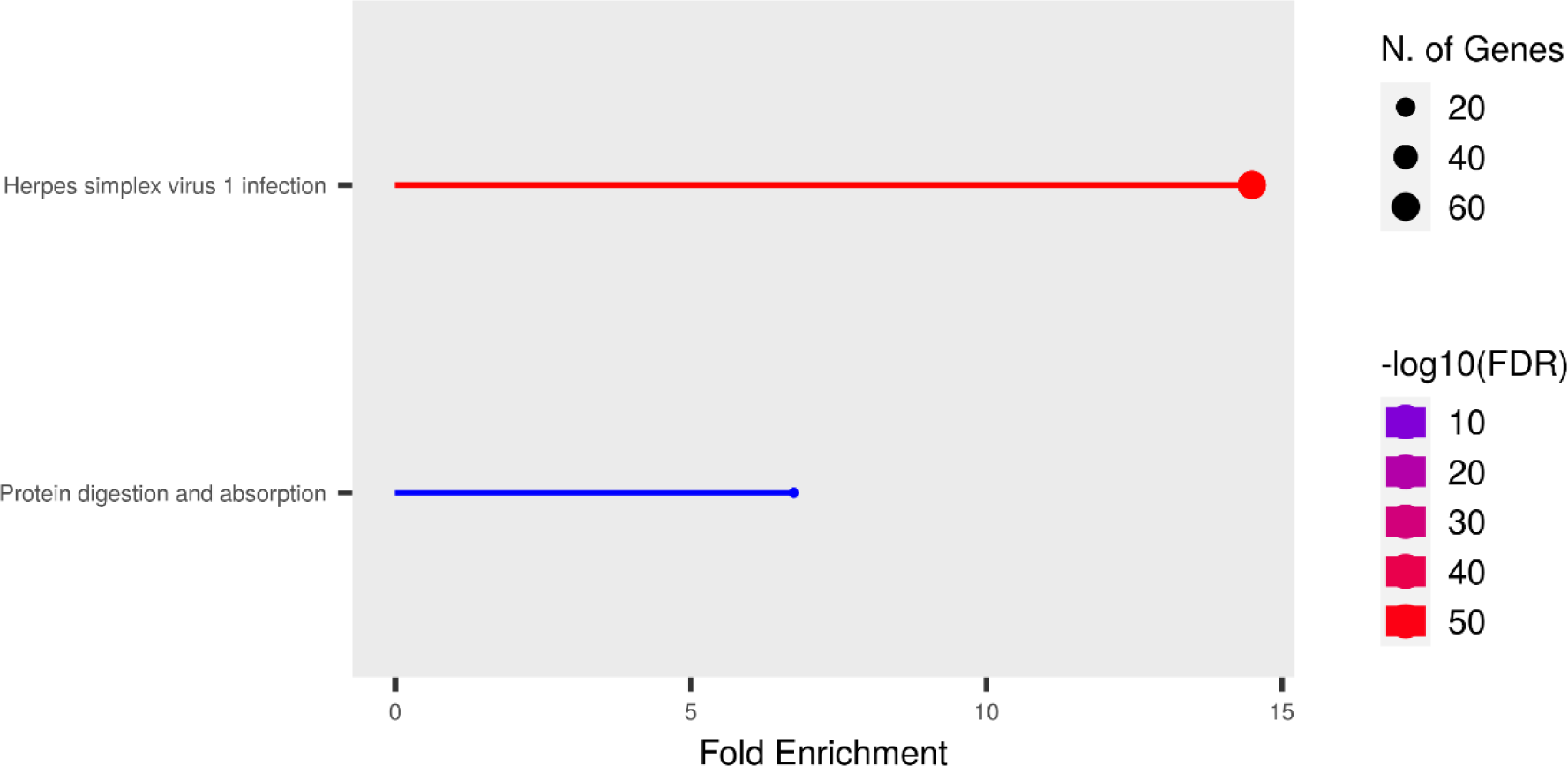
Gene enrichment analysis of KEGG pathways on bottom-right corner in human APOBEC3A/B TC motif CDUR plot of gene.

**Figure S4.**
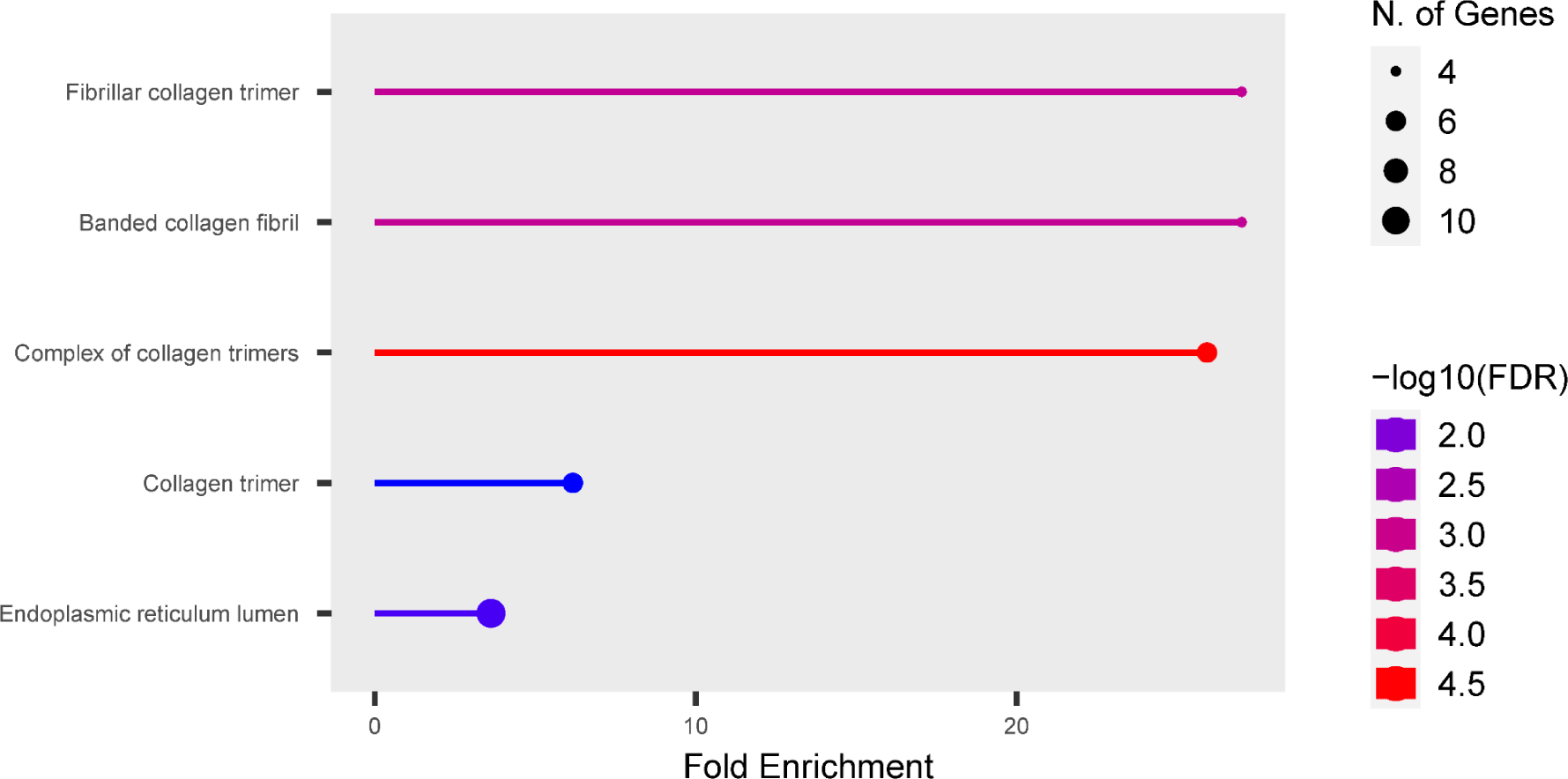
Gene enrichment analysis of GO Cellular Component on bottom-right corner in human APOBEC3A/B TC motif CDUR plot of gene.

**Figure S5.**
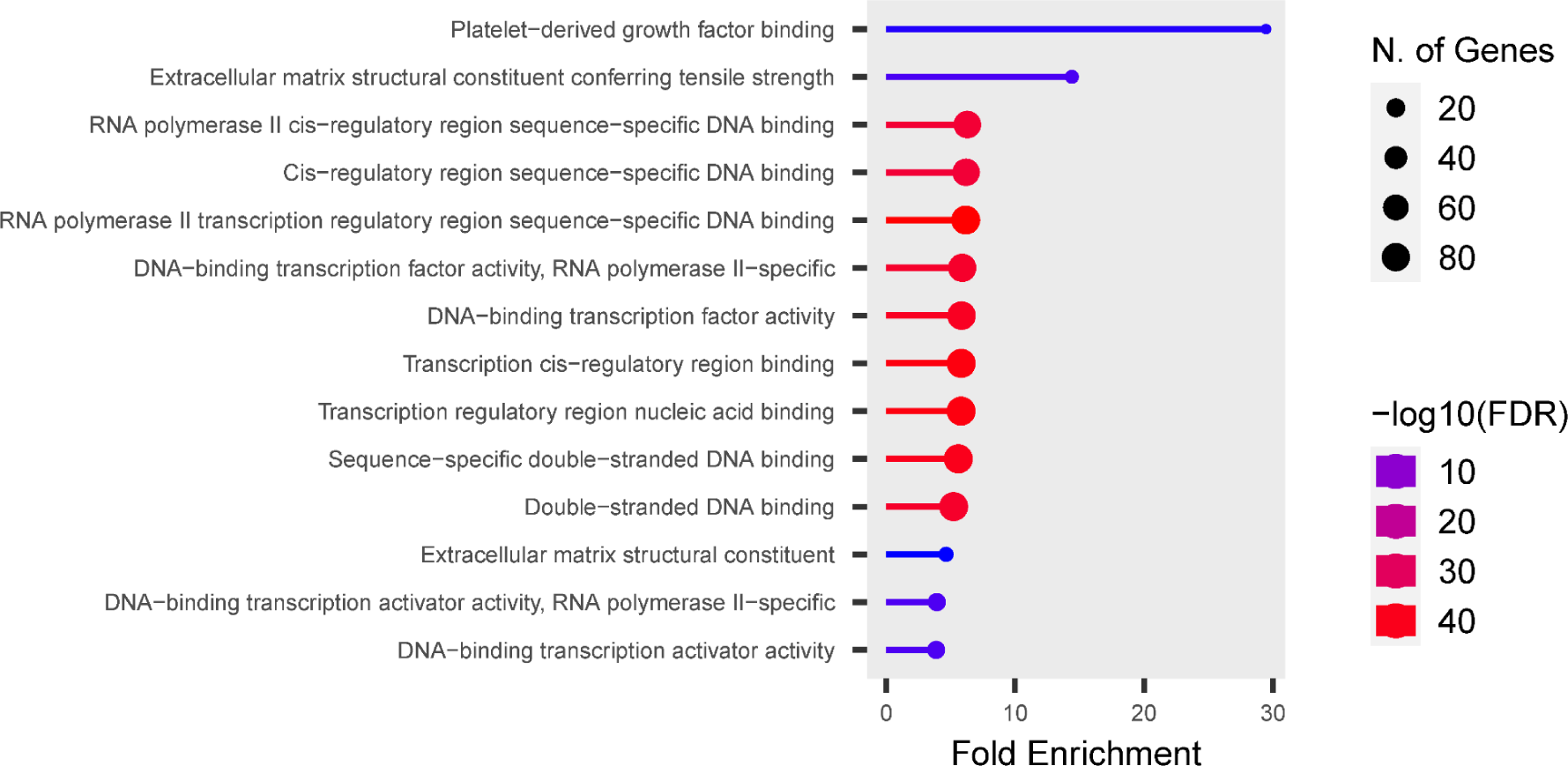
Gene enrichment analysis of GO Molecular Functions on bottom-right corner in human APOBEC3A/B TC motif CDUR plot of gene.

**Figure S6.**
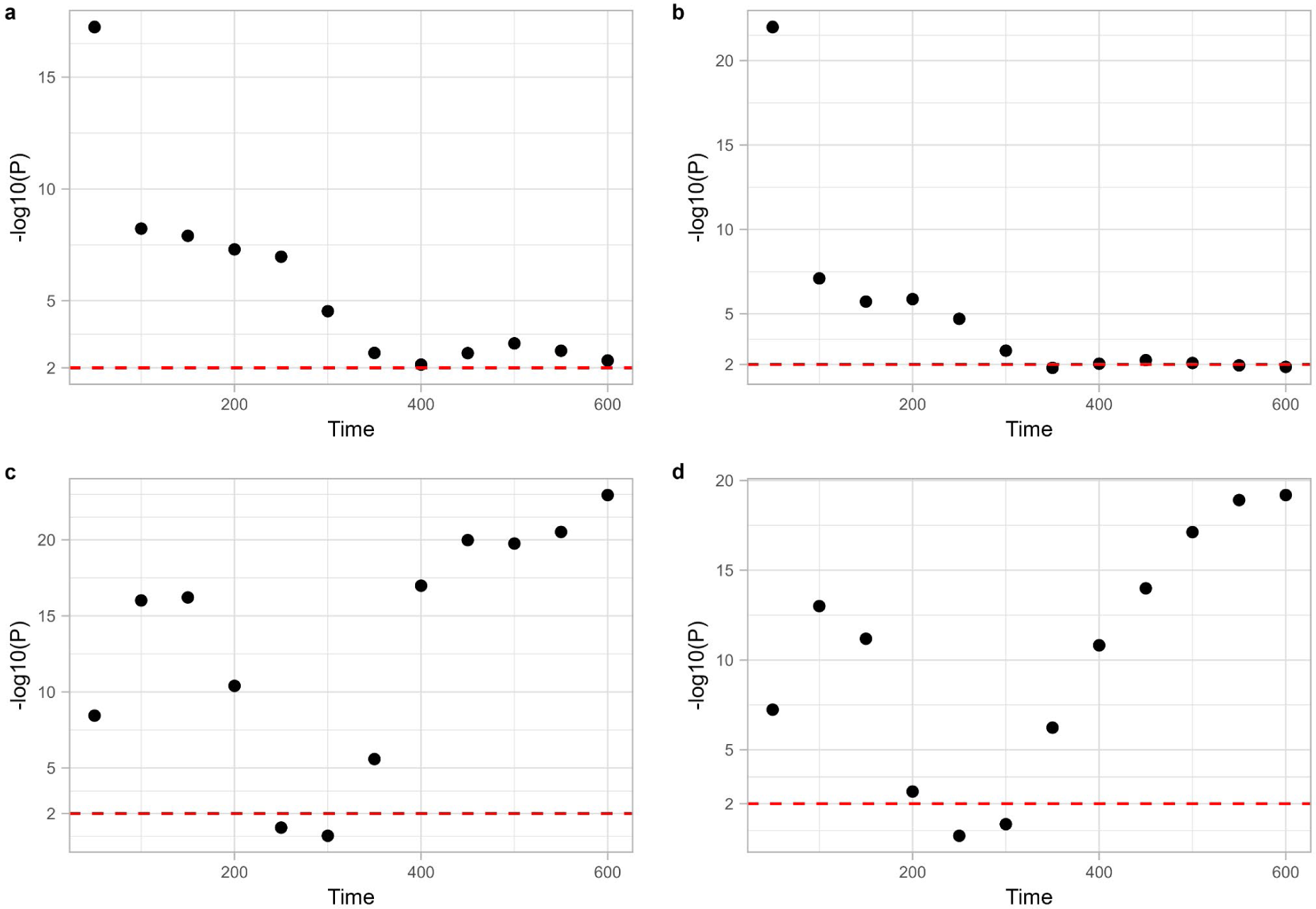
Statistical significance between simulations at every 50 time points by the number of genotypes and heterogeneity. P-values were obtained by t-test on each simulation at every 50 time points. a. Statistical difference by the number of genotypes between with and without APOBEC mutations decreases over time. b. Statistical difference by heterogeneity between with and without APOBEC mutations decreases over time. c. tatistical difference by the number of genotypes between uniform and bimodal distribution increase over time. d. Statistical difference by the number of genotypes between uniform and bimodal distribution increase over time.

**Figure S7.**
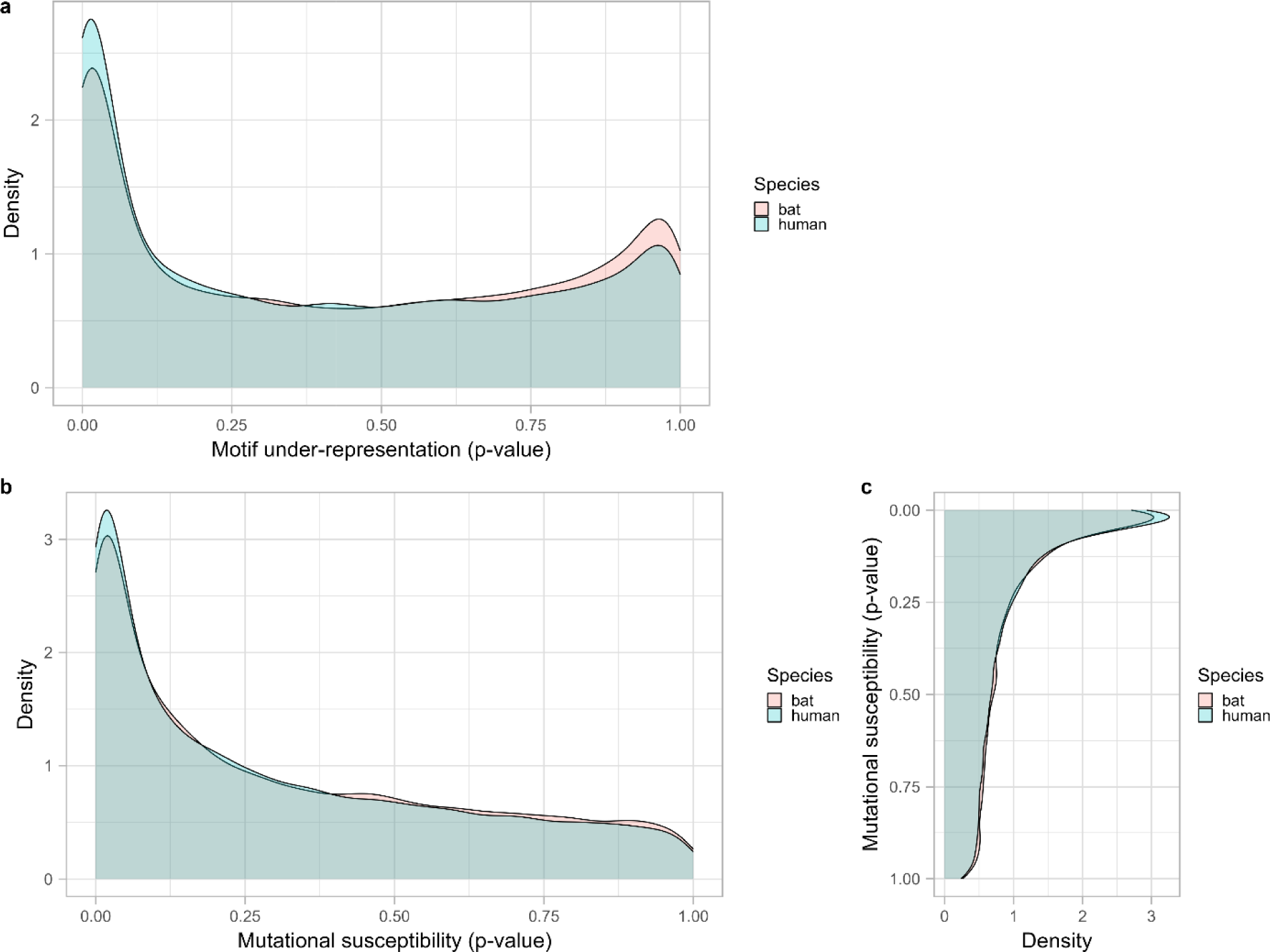
Comparison between human and *Pteropus alecto* genome motif under-representation and mutational susceptibility.

**Figure S8.**
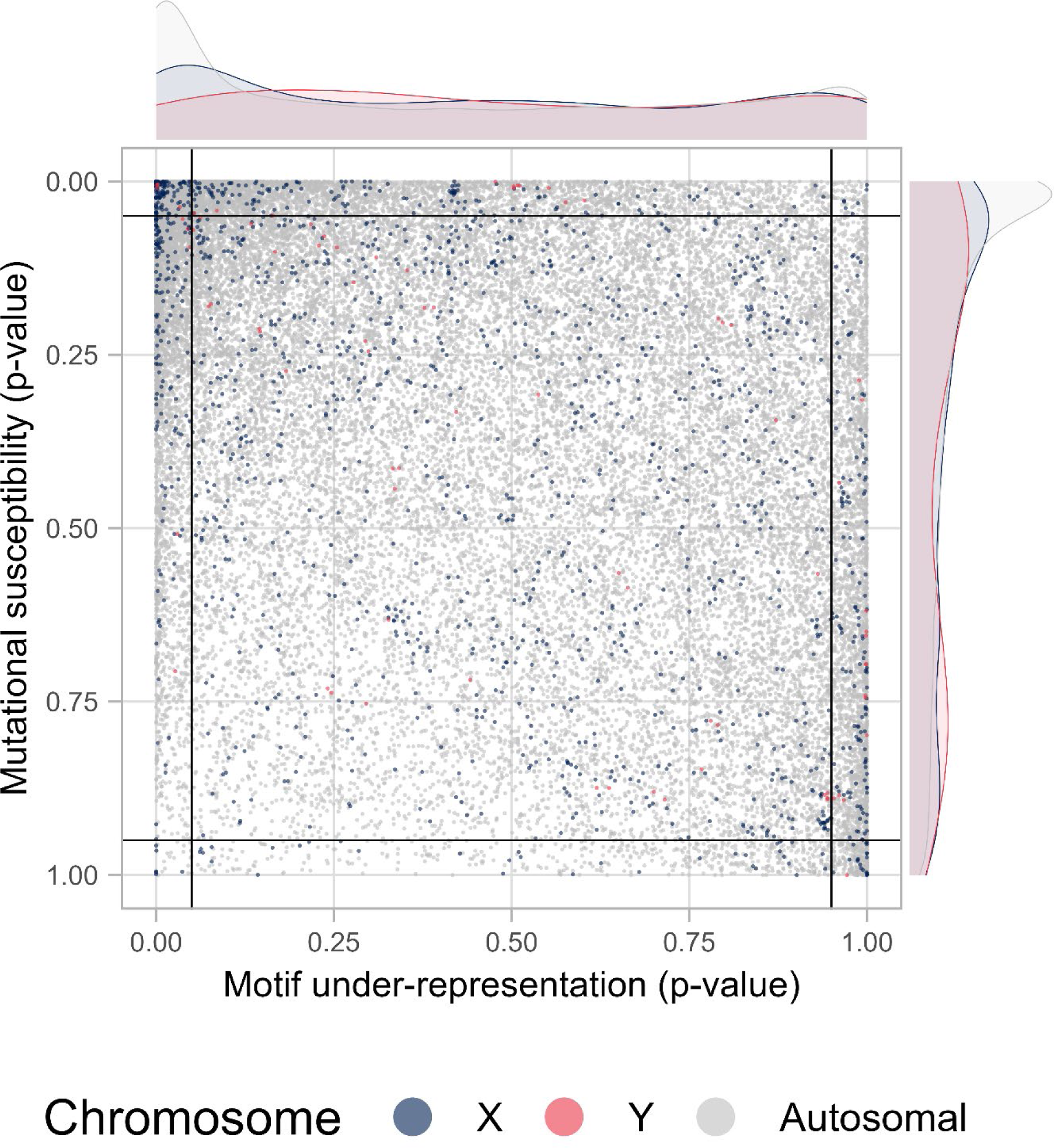
Distribution comparison between genes from two sex chromosome and autosomal chromosome.

**Figure S9.**
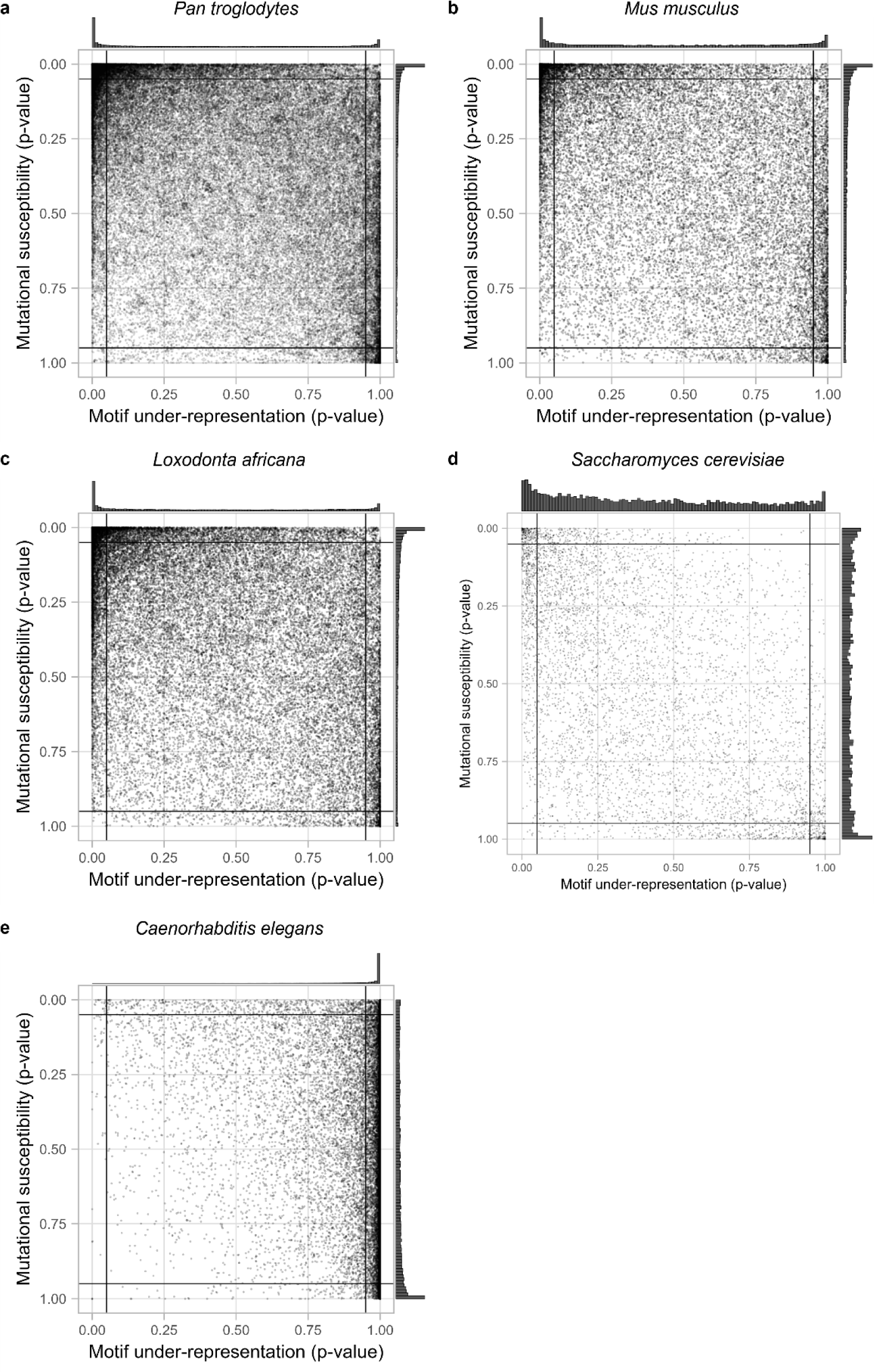
CDUR plots for 5 organisms: Chimpanzee, mice, elephant, yeast, and *C. elegans*.

**Figure S10.**
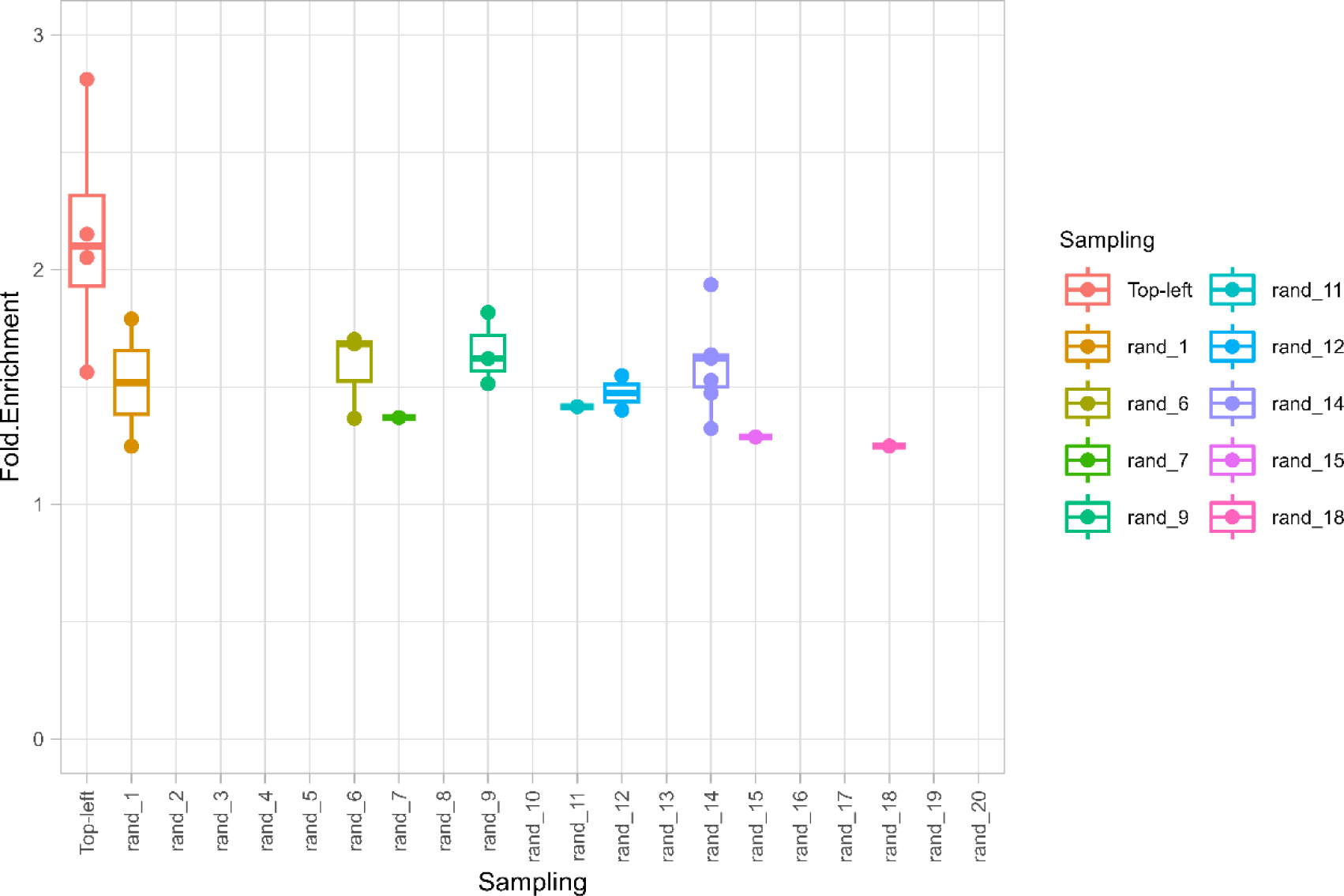
Human top-left corner genes KEGG pathway enrichment compared to the same number of randomly selected genes.

