## Supplementary material for "Evolvability of cancer-associated genes under APOBEC3A/B selection": Suuplementary Figures

### Supplemental information

### Supplementary figures

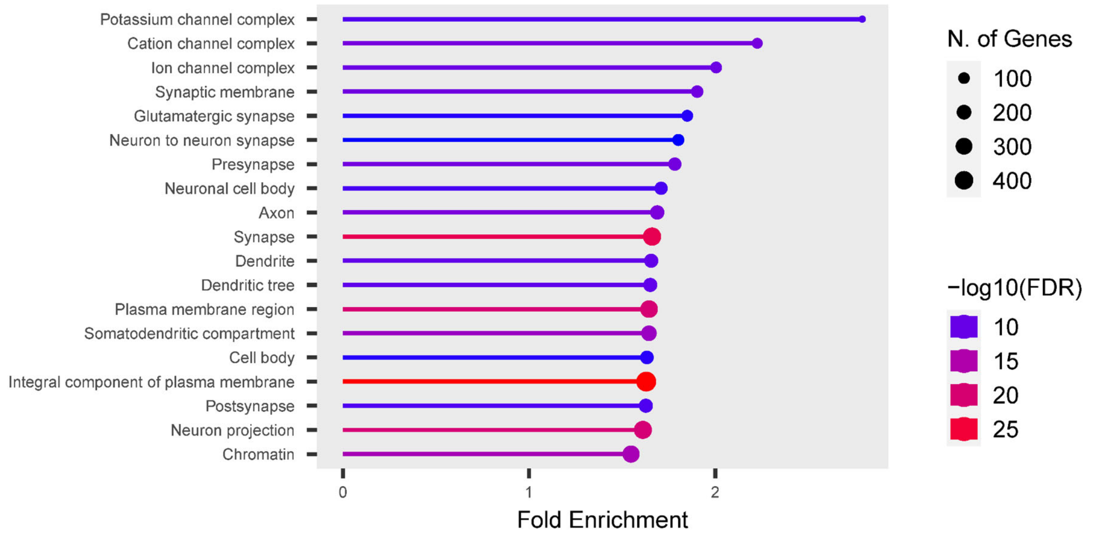


**Figure S1. Gene enrichment analysis of GO Cellular Component on top-left corner in human APOBEC3A/B TC motif CDUR plot of gene.**

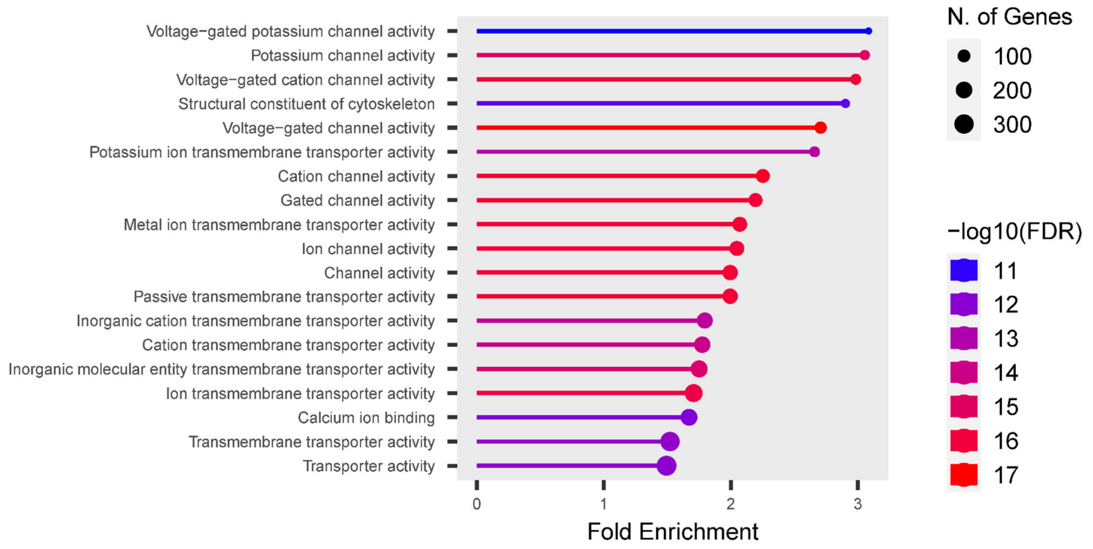


**Figure S2. Gene enrichment analysis of GO Molecular Functions on top-left corner in human APOBEC3A/B TC motif CDUR plot of gene.**

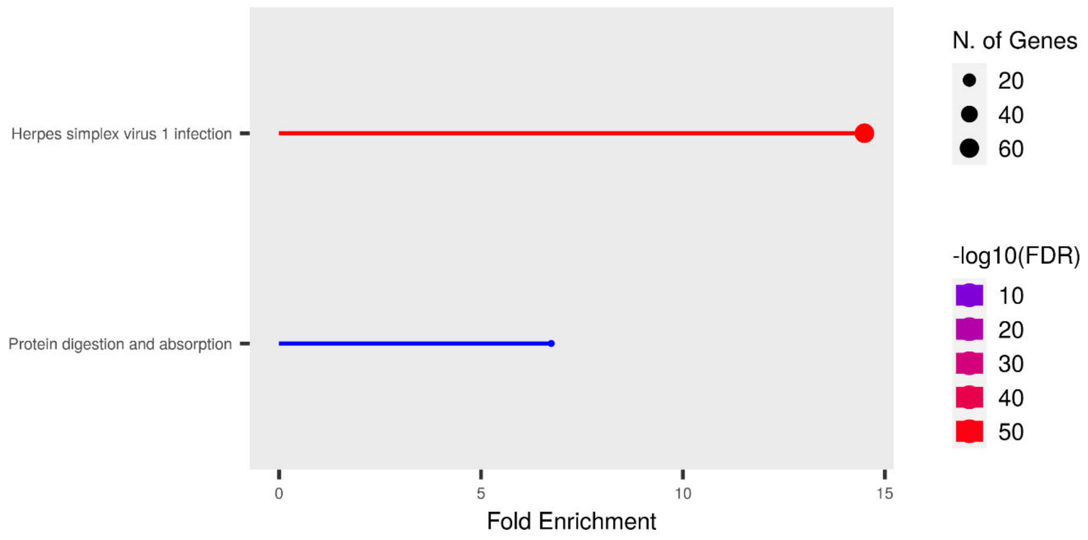


**Figure S3. Gene enrichment analysis of KEGG pathways on bottom-right corner in human APOBEC3A/B TC motif CDUR plot of gene.**

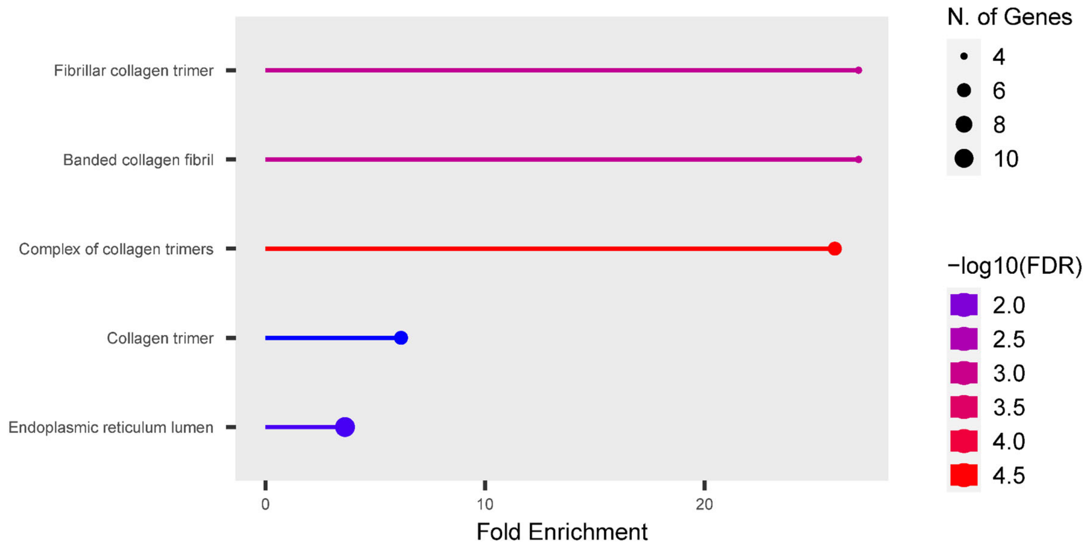


**Figure S4. Gene enrichment analysis of GO Cellular Component on bottom-right corner in human APOBEC3A/B TC motif CDUR plot of gene.**

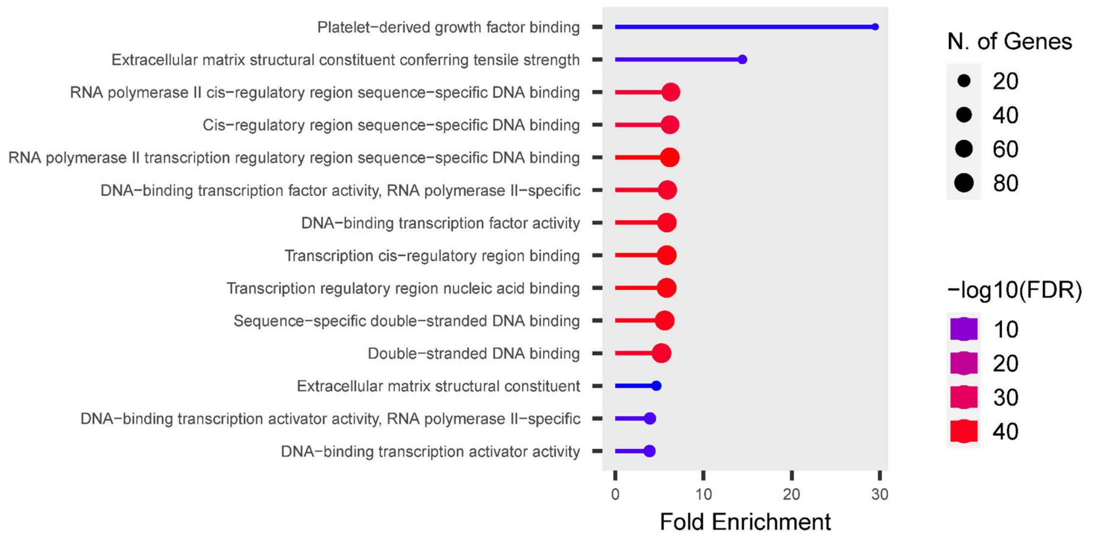


**Figure S5. Gene enrichment analysis of GO Molecular Functions on bottom-right corner in human APOBEC3A/B TC motif CDUR plot of gene.**

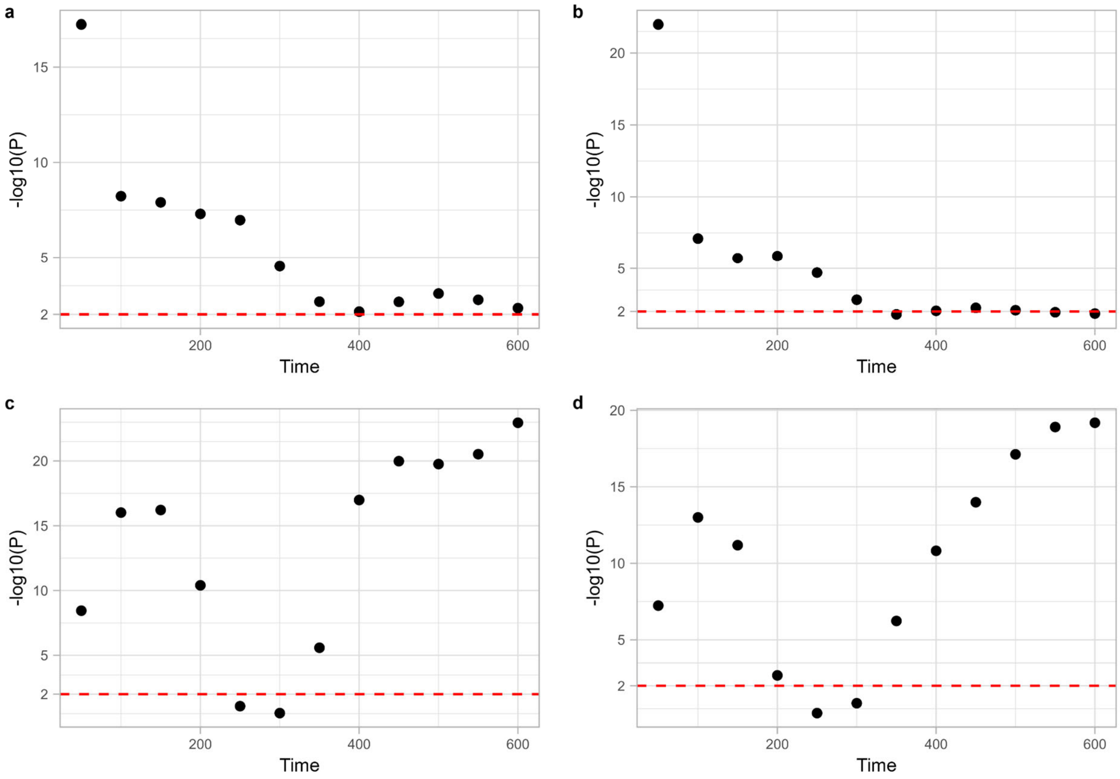


**Figure S6. Statistical significance between simulations at every 50 time points by the number of genotypes and heterogeneity.** P-values were obtained by t-test on each simulation at every 50 time points. a. Statistical difference by the number of genotypes between with and without APOBEC mutations decreases over time. b. Statistical difference by heterogeneity between with and without APOBEC mutations decreases over time. c. Statistical difference by the number of genotypes between uniform and bimodal distribution increase over time. d.  Statistical difference by the number of genotypes between uniform and bimodal distribution increase over time.

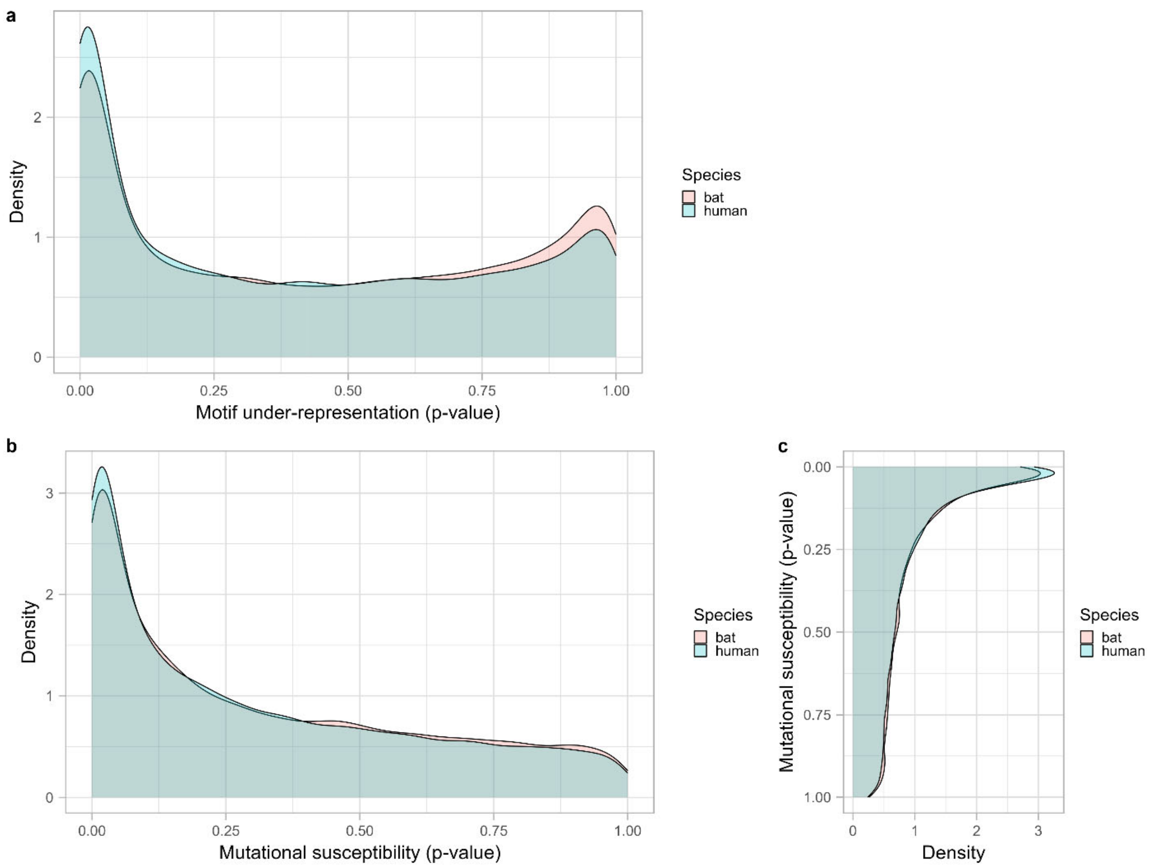


**Figure S7. Comparison between human and *Pteropus alecto* genome motif under-representation and mutational susceptibility.**

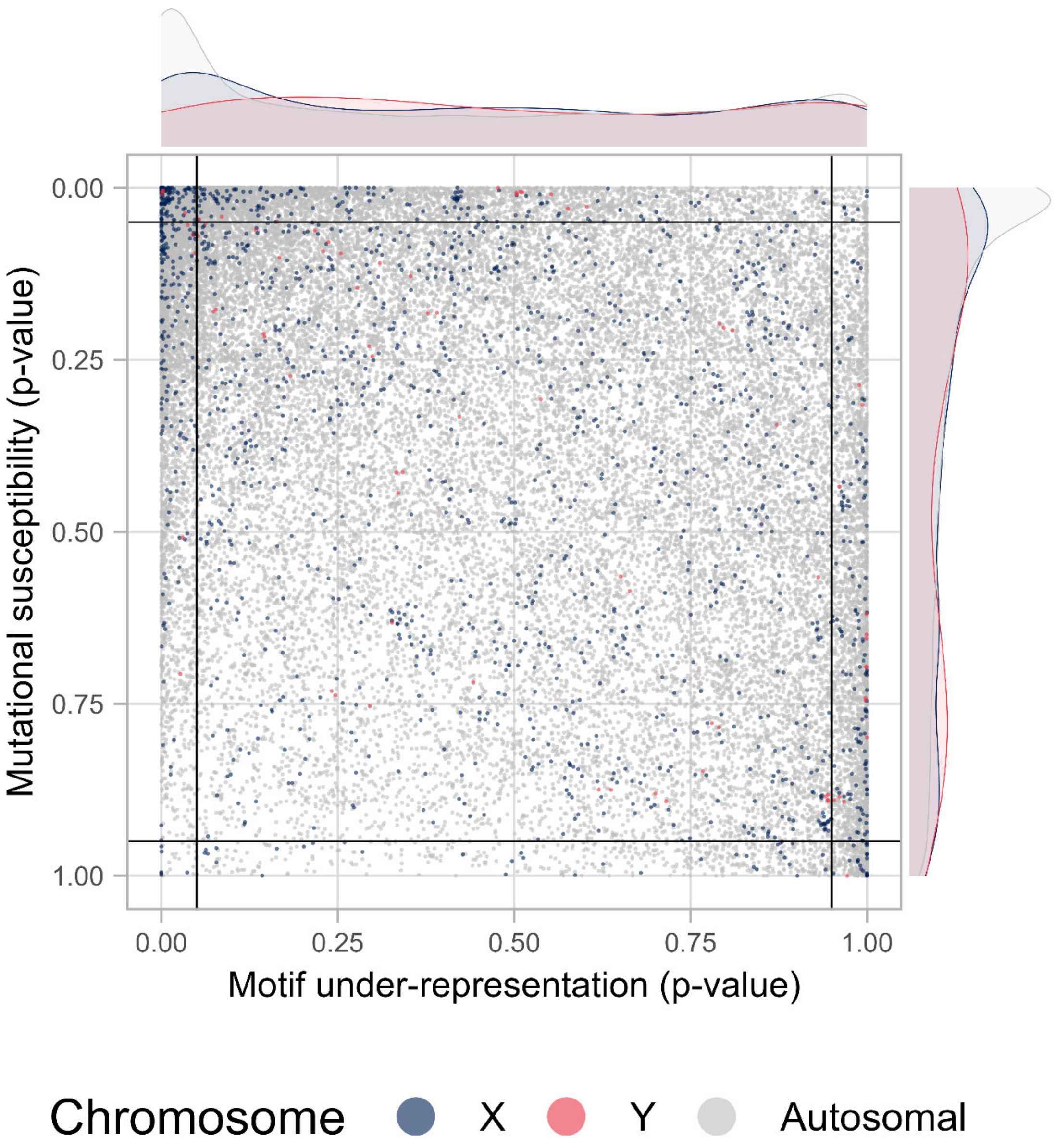


**Figure S8. Distribution comparison between genes from two sex chromosome and autosomal chromosome.**

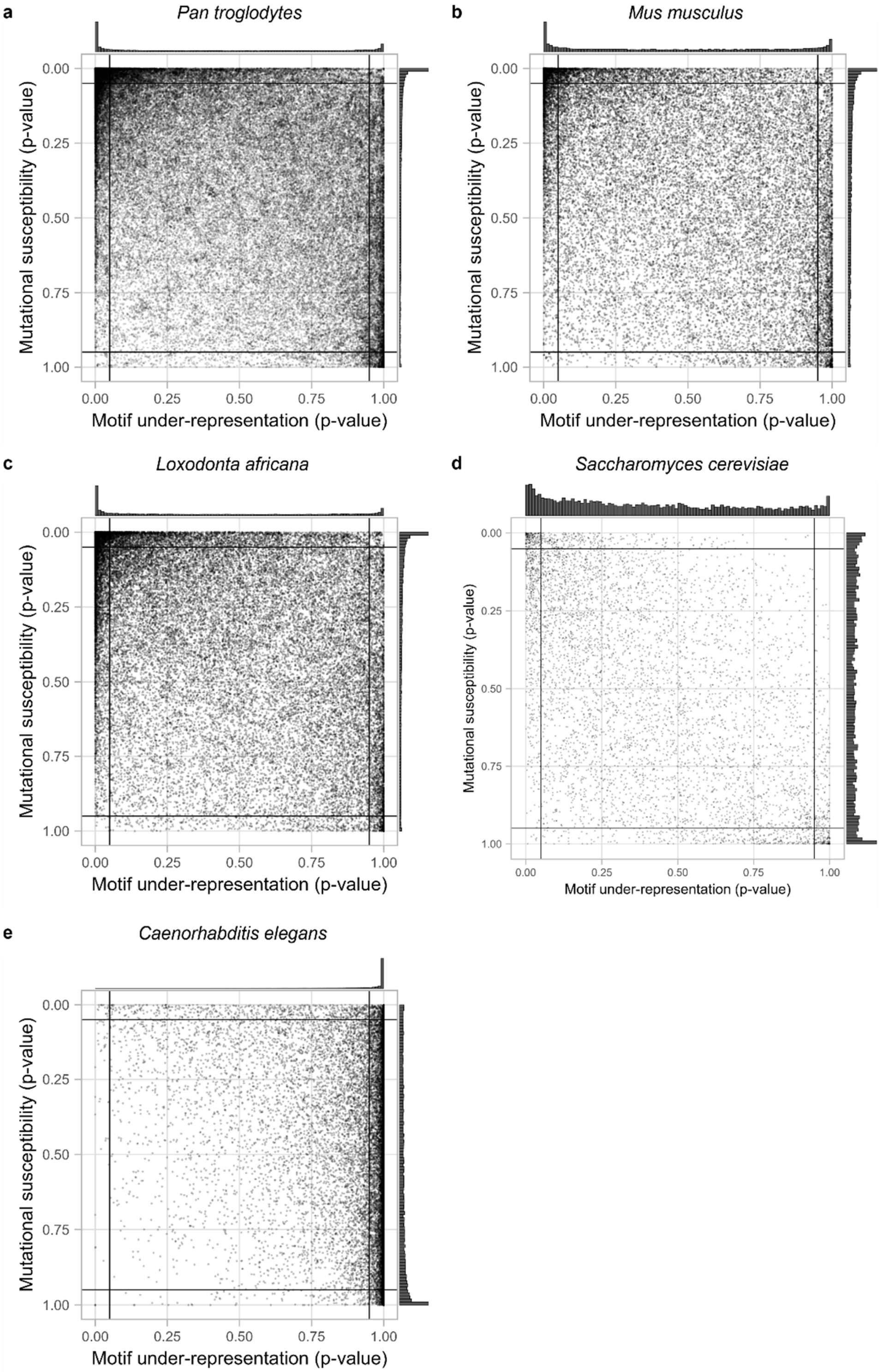


**Figure S9. CDUR plots for 5 organisms: Chimpanzee, mice, elephant, yeast, and *C. elegans*.**

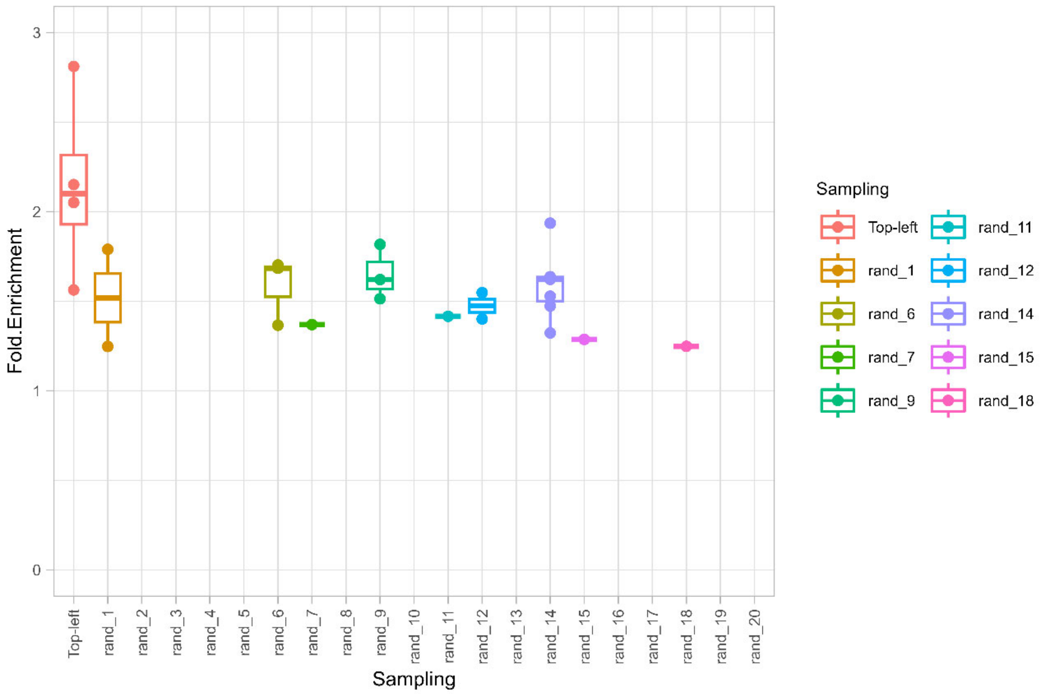


**Figure S10. Human top-left corner genes KEGG pathway enrichment compared to the same number of randomly selected genes.**
